# High-resolution promoter interaction analysis implicates genes involved in the activation of Type 3 Innate Lymphoid Cells in autoimmune disease risk

**DOI:** 10.1101/2022.10.19.512842

**Authors:** Valeriya Malysheva, Helen Ray-Jones, Nora Lakes, Rachel A. Brown, Tareian A. Cazares, Owen Clay, David E. Ohayon, Pavel Artemov, Joseph A. Wayman, Zi F. Yang, Monica Della Rosa, Carmen Petitjean, Clarissa Booth, Joseph I.J. Ellaway, Jenna R. Barnes, Andrew W. Dangel, Ankita Saini, William R. Orchard, Xiaoting Chen, Sreeja Parameswaran, Frances Burden, Mattia Frontini, Takashi Nagano, Peter Fraser, Stefan Schoenfelder, Matthew T. Weirauch, Leah C. Kottyan, David F. Smith, Nick Powell, Jill M. Weimer, Eugene M. Oltz, Chris Wallace, Emily R. Miraldi, Stephen Waggoner, Mikhail Spivakov

## Abstract

Innate lymphoid cells (ILCs) are rare, tissue-resident innate lymphocytes that functionally mirror CD4+ T helper cell lineages but lack antigen receptors. Type 3 ILCs (ILC3s) are enriched in the gut, airways, and mucosal lymphoid tissues, where they regulate inflammation and promote barrier integrity. To define the regulatory architecture of primary human ILC3s, we map promoter-anchored chromosomal contacts using high-resolution, low-input Promoter Capture Hi-C (PCHi-C) in these cells alongside CD4+ T cells. By combining statistical detection with a PCHi-C-adapted Activity-by-Contact approach, we link promoters to distal regulatory elements, identifying hundreds of ILC3-specific contacts. We use these maps to connect genome-wide association study (GWAS) risk variants for Crohn’s disease to target genes using multiCOGS, a Bayesian framework that integrates PCHi-C with summary-statistic imputation and multivariate fine-mapping. This analysis highlights both known and unanticipated candidates, including *CLN3*, a causal gene for the neurodevelopmental Batten disease. Using a mouse ILC3-like cell line, we show that *Cln3* is downregulated upon cytokine stimulation, and *Cln3* overexpression alters stimulation-induced transcriptional programmes and cytokine secretion. Extending this approach, we generate a catalogue of ILC3-linked risk genes for five additional autoimmune conditions and show that they are enriched for regulators of the ILC3 inflammatory response identified in a CRISPR interference screen. Together, these findings illuminate long-range gene control in ILC3s and prioritise known and newly implicated autoimmune risk genes with potential roles in this clinically important cell type.

## Introduction

Innate lymphoid cells (ILCs) play crucial roles in inflammation and immunity, as well as in tissue development and homeostasis^1,2^. ILCs develop from common lymphoid progenitors and share many features with CD4+ T lymphocytes, but do not express rearranged T cell receptors^3^. Therefore, rather than acting as part of the adaptive immune system, ILCs respond to cytokines and pathogens from the environment by producing regulatory cytokines and exerting immunomodulatory activity^4,5^.

Three main types of ILCs have been identified based on their cytokine profiles and the transcription factors regulating their development and function^2,3^. The first group includes tissue-resident ILC1s that play a role in immune defence against viruses and certain bacteria^6,7^. The second group consists of ILC2s, which regulate airway and skin inflammatory responses and are implicated in disorders such as asthma and atopic dermatitis^6^. Finally, the third group includes lymphoid tissue-inducer cells, which are involved in lymph node development, and ILC3s, which participate in host defence and the maintenance of epithelial barrier homeostasis^2–4^. The ILC3 population is distributed across multiple tissues, including the gut, where they are essential for mucosal homeostasis and barrier integrity^8^. ILC3-derived cytokines such as IL-17 and IL-22 promote epithelial cell renewal and release of antimicrobial peptides^9^. However, overexpression of these cytokines in the gut has been associated with the development or exacerbation of Crohn’s disease (CD)^10–12^.

Immune disorders, including CD, are known to have a significant genetic component, with genome-wide association studies (GWAS) identifying hundreds of disease susceptibility variants associated with these conditions^13^. Given the importance of ILCs in immune control, it is highly plausible that some of these variants affect ILC function. However, as most GWAS variants are non-coding and these studies are, by design, cell-type agnostic, identifying causal genes and cell types implicated by GWAS variants is often challenging.

GWAS variants are strongly enriched at transcriptional enhancers^14–16^, and therefore, cell type-specific maps of active enhancers and enhancer-promoter connections provide important clues for the functional interpretation of GWAS findings^17,18^. Recent studies have mapped ILC enhancers by the assay for transposase-accessible chromatin (ATAC-seq) and chromatin immunoprecipitation (ChIP-seq) for the H3K27ac histone mark, identifying putative key regulators of ILC identity and their downstream targets based on proximal gene assignment^19–23^. However, enhancers often localise large distances (up to megabases) away from their target gene promoters, physically contacting them in the 3D space of the nucleus in a cell-type-specific manner. Therefore, robust and sensitive identification of enhancer-promoter contacts, which is instrumental for inferring the effector genes of non-coding GWAS variants, requires robust and sensitive profiling of chromosomal architecture.

Chromosome conformation capture assays such as Hi-C, which are based on the proximity ligation of cross-linked, digested chromatin, provide powerful tools for connecting enhancers and GWAS variants with target genes^24,25^. The conventional Hi-C technique theoretically allows the detection of all pairwise chromosomal contacts across the genome. However, the complexity of the resulting sequencing libraries requires extremely high sequencing coverage to achieve the sensitivity and resolution needed for the detection of specific enhancer-promoter contacts. This challenge can be addressed by techniques such as Capture Hi-C that selectively enrich Hi-C material for contacts involving, at one end, regions of interest such as gene promoters^26–29^. Over the last decade, we and others have demonstrated the power of Promoter Capture Hi-C (PCHi-C) in determining transcriptional regulatory circuitries and in linking enhancers and disease-associated genetic variants with putative target genes^30–36^. In foundational studies^30,31^, we applied this approach to 17 abundant human primary blood cell types and developed COGS (Capture Hi-C Omnibus Gene Score), a Bayesian approach for prioritisation of GWAS target genes using statistical fine-mapping and PCHi-C data. Results from this work were incorporated into major variant-to-gene resources, including OpenTargets Genetics^37^ and Priority Index^38^. However, the PCHi-C protocol used in these studies required dozens of millions of input cells, precluding the analysis of rare cell types.

Here, we address this limitation by using a high-resolution and efficient PCHi-C protocol to profile the *cis*-regulatory wiring of ILC3s isolated from human tonsils^30^. We detect promoter-enhancer contacts in PCHi-C data using a combination of our established statistical interaction-calling methodology (CHiCAGO)^39,40^ and a newly developed adaptation of the Activity-by-Contact^14,41^ (ABC) approach to PCHi-C data that we term Activity-by-Captured-Contact (ABCC). We develop a modified PCHi-C-aware GWAS gene prioritisation algorithm, multiCOGS, that incorporates summary statistics imputation and multivariate statistical fine-mapping, and use it to prioritise known and novel genes for CD through chromatin contacts. Several of the genes are uniquely prioritised using PCHi-C data from ILC3s but not CD4+ T cells, including the *CLN3* gene, mutations in which underpin ∼80% of cases of the neurodegenerative disorder Batten disease^42,43^. We show that this gene is downregulated upon cytokine stimulation of mouse ILC3s, and *Cln3* overexpression in an ILC3-like mouse cell line influences stimulation-responsive transcriptional programmes and cytokine production. Finally, expanding multiCOGS to five additional autoimmune conditions, we generate a catalogue of effector genes implicating ILC3s and show that they are enriched among putative regulators of ILC3 inflammatory function. Together, our results shed light on ILC3 cis-regulatory circuitries and prioritise autoimmune risk effector genes with potential roles in this clinically important cell type.

## Results

### A compendium of promoter-anchored chromosomal contacts in human ILC3s

To profile promoter-anchored chromosomal contacts in type 3 innate lymphoid cells (ILC3s), we employed our low-input *DpnII*-based PCHi-C protocol^44,45^ on ILC3s extracted from human tonsils (**Fig. 1A**). Significant promoter contacts were detected with CHiCAGO^39^ at a single-fragment resolution, as well as after pooling the ‘other end’ fragments into ∼5 kb bins, while leaving the baited promoter-containing fragment unbinned (Methods)^40^. Using this approach, we detected 31,003 contacts between promoters and promoter-interacting regions (PIRs) at a single-fragment resolution and 58,632 contacts in 5 kb bins (**Fig. 1B**; **Table S1**; **Data S1-S2** at https://osf.io/aq9fb). Binning resulted in the detection of longer-range contacts, as we reported previously in other cell types^40^ (**Fig. 1C, D**). A joint dimensionality reduction analysis^46^ of ILC3 promoter interaction profiles with those detected in 17 abundant blood cell types using *HindIII-*based PCHi-C segregated ILC3s with other lymphoid cell types, consistent with the notion that patterns of promoter interactions reflect the cells’ lineage history^30^ (**Fig. S1A**; see Methods).

**Figure 1.**
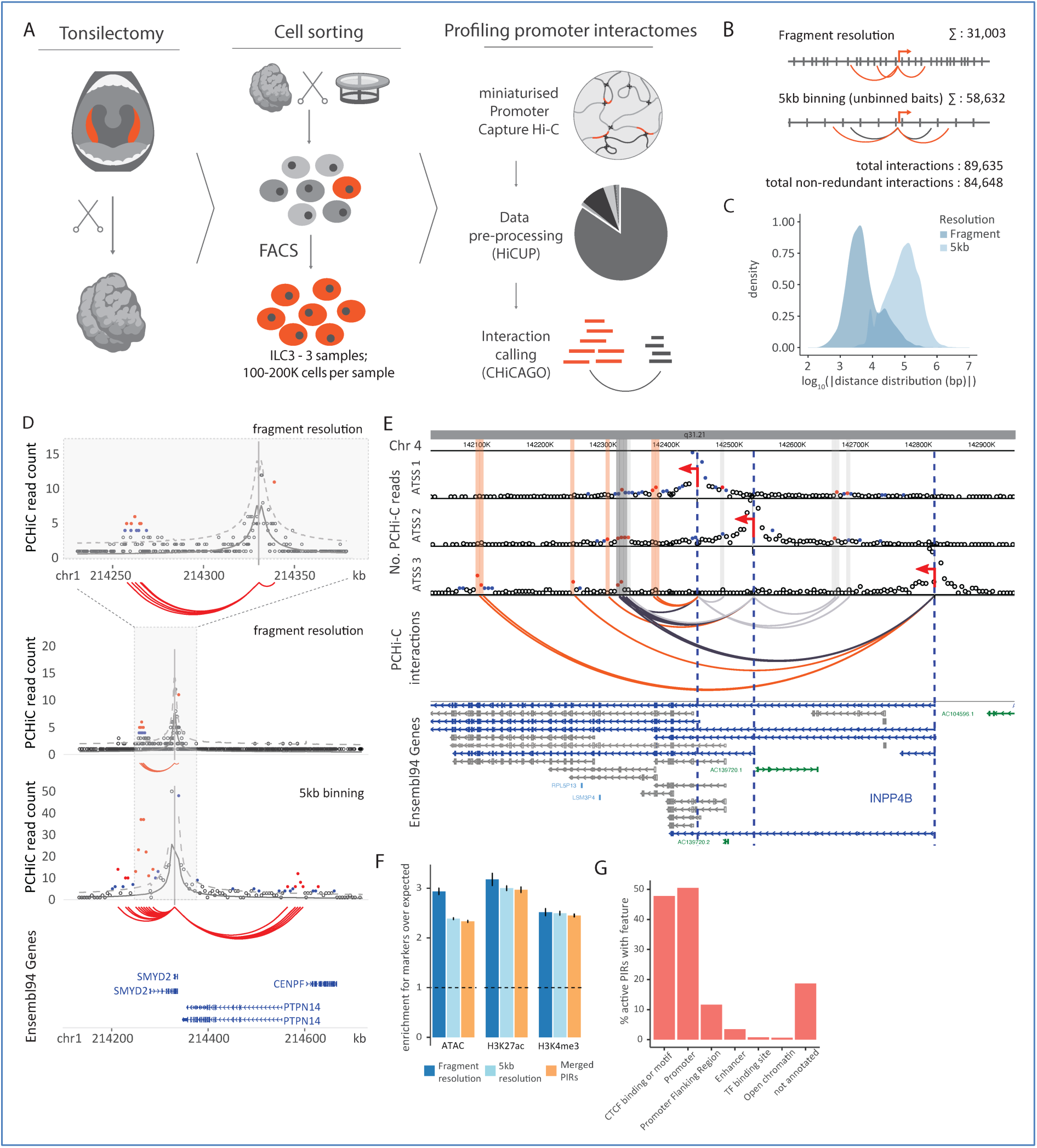
Compendium of promoter interactions in ILC3s. **A.** Outline of the study. **B.** Chromosomal interaction binning strategy. The analysis is done in two modes: fragment resolution (no binning) and 5kb binning. In the 5kb binning mode, the baited (captured) fragment containing a gene promoter, is left unbinned to enable high-resolution linkage between the promoter and distal enhancers. Interactions uniquely detected in one mode only are shown as red arcs, and those detected in both modes are shown as grey arcs. The numbers of significant interactions are given for each mode individually and merged across both modes (see Methods for details). **C.** Comparison of promoter-PIR distance distributions for PIRs detected at fragment and 5kb resolution. **D.** Example of chromosomal interactions for the *SMYD2* gene at fragment and 5kb resolution. The inset shows a zoomed-in view of the promoter interactions detected at fragment resolution. **E.** Example of multiple degrees of contact sharing between alternative promoters for the *INPP4B* gene. Captured alternative promoters are indicated by red arrows and blue dashed lines. The transcripts driven by these promoters (based on Ensembl 94) are shown in blue, and other *INPP4B* transcripts are shown in grey. Transcripts for processed pseudogenes are shown in light blue and lincRNAs in green. PIRs are categorised as fully shared between alternative promoters (dark grey arcs), partially shared (light grey arcs) or distinct (red arcs). **F.** Enrichment of PIRs for the markers of active enhancers and promoters (H3K27ac and H3K4me3) and accessible chromatin (ATAC) in hILC3s. The error bars represent 95% confidence intervals, accounting for error propagation. **G.** Characterisation of active and/or open ILC3 PIRs at merged fragments as per Ensembl annotations and CTCF motifs.

The increased resolution afforded by using *DpnII* in Hi-C library generation enabled capturing alternative transcription start sites (ATSSs) for 6,789 genes located on separate *DpnII* fragments. Remarkably, genes with captured ATSSs displayed distinct interaction landscapes across isoforms (**Fig. S1B, C, D**). The three ATSSs of the *INPP4B* gene provide examples of the multiple degrees of contact sharing across its 14 PIRs included in the analysis (**Fig. 1E**).

Next, we explored the epigenetic status of detected PIRs and compared the chromatin profile of ILC3s with those of 88 other blood cell types detected by the Ensembl regulatory build^47^. As expected, at both fragment and 5-kb resolution ILC3 PIRs were enriched for markers of accessible and/or active enhancers (ATAC, H3K27ac) and active transcription (H3K4me3), based on public data in this cell type isolated from tonsils of pediatric donors^21^ (“active PIRs”, **Fig. 1F**). Nearly half of all accessible and/or active ILC3 PIRs (47.8%, 8,718/18,231) overlapped with annotated CTCF motifs or CTCF binding events in at least one of the Ensembl-profiled cell types (**Fig. 1G**), consistent with the key role of CTCF in 3D chromosomal organisation. However, only 3% of active/open regions in ILC3s (636/18,231) contained Ensembl enhancer annotations^48^, while nearly 20% of accessible and/or active PIRs (3,411/18,231) did not have any functional annotations in the Ensembl data (**Fig. 1G**).

We then considered the overlap of the active and/or accessible PIRs in ILC3s with those in 17 abundant blood cell types profiled with PCHi-C at *HindIII* resolution^30^. In contrast to chromatin annotations, the majority of active/accessible PIRs in ILC3s also had promoter contacts in these blood cell types (∼80.4%, 12,409/15,435). Furthermore, ∼60% of the active PIRs (9,054/15,435) contacted the same gene promoters in both ILC3s and other blood cells **(Data S3** at https://osf.io/aq9fb). Consistent with previous observations, this result confirms that patterns of promoter-enhancer contacts are more preserved across related lineages compared with enhancer activity *in cis*^49^. We then probed the relationship between enhancer-promoter connectivity and gene expression. For this, we integrated promoter-enhancer interactions detected here with publicly available single-cell gene expression data (scRNA-seq) in human mucosal tissue ILC3s^50^. In agreement with epigenetic studies in other cell types,^30^ we observed a significant positive correlation between the number of active and/or open PIRs and gene expression (**Fig. S1E**).

Overall, our analysis provides a high-resolution compendium of promoter contacts in ILC3s, including novel ILC3-specific regulatory elements and divergent contacts at ATSSs.

### Inference of enhancer-promoter interactions using Activity-by-Captured-Contact (ABCC) complements significant interaction detection

To further increase the sensitivity of detecting functional promoter-enhancer chromosomal interactions from PCHi-C data, we adapted the Activity-by-Contact (ABC) approach^41^ originally developed for Hi-C. In contrast to CHiCAGO, which detects significant interactions relative to a distance-dependent background, ABC considers any observed contact frequency between a chromatin region and a promoter as potentially functionally meaningful, irrespective of whether this frequency exceeds that expected by chance. In addition, while CHiCAGO scores are independent of enhancer activity levels at the PIRs, ABC incorporates both contact frequency and enhancer activity into the final metric (“ABC score”)^41^.

In our adaptation of ABC, which we term ‘Activity-by-Captured Contact’ (ABCC), we estimated contact frequencies from imputed PCHi-C data, leveraging the statistical modelling of these data produced by CHiCAGO for the imputation task (**Fig. 2A**, **Fig. S2A, S2B**, see Methods). To validate the ability of the ABCC algorithm to detect functional enhancer-promoter pairs, we took advantage of CRISPR interference (CRISPRi) enhancer perturbation data in K562 cells, which was generated to validate the original ABC approach^14^. As inputs for ABCC, we used public epigenetic annotations in K562 cells and our previously generated high-coverage PCHi-C data in their physiological counterparts, erythroblasts^30^. These analyses demonstrated the power of ABCC to predict functional enhancer-promoter links from lineage-relevant PCHi-C and chromatin readouts (**Fig. S2C**). In contrast, using PCHi-C data from lymphoid cells at an equivalent coverage reduced ABCC performance (**Fig. S2C)**. In addition, joint clustering of the ABCC profiles generated for four primary blood cell types successfully reconstructed the lineage relationships between them (**Fig. S2D**). These results highlighted the potential of ABCC to infer lineage-specific *cis-*regulatory architecture. In comparison with CHiCAGO, ABCC generally detected shorter-range promoter interactions, which was expected due to its reliance on raw contact frequencies (**Fig. S2E**). Both ABCC- and CHiCAGO-detected contacts were enriched for markers of accessible (DNase-seq) and/or active (H3K27ac) enhancers, with regions called by both approaches showing the highest enrichment for these marks (**Fig. S2F**). Taken together, these results suggest that ABCC and CHiCAGO detect complementary subsets of regulatory promoter contacts.

**Figure 2.**
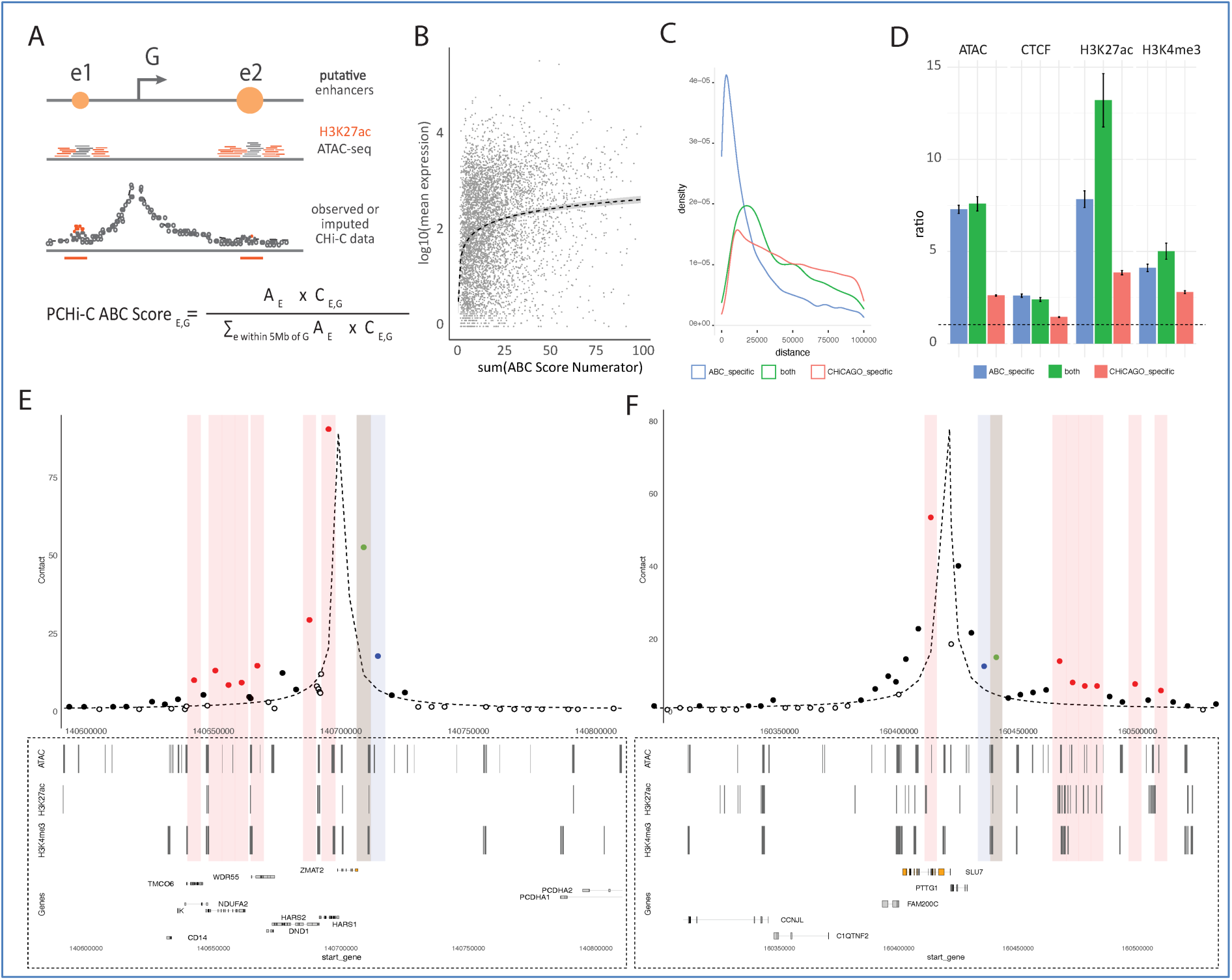
Combining ABCC and CHiCAGO to link distal elements with target genes. **A.** Schematic depicting the adaptation of the Activity-By-Contact (ABC) model for use with PCHi-C data, termed Activity-By-Captured-Contact (ABCC). **B.** Correlation between gene expression and ABC numerator score summed across all predicted enhancers per gene. The dashed line shows a mixed model fit via restricted maximum likelihood, with the shaded area around the line representing the confidence interval. **C.** Interaction distance comparison across CHiCAGO-specific, ABCC-specific and shared interactions. **D.** Enrichment for markers of active/open regulatory elements in CHiCAGO-specific, ABCC-specific, and shared regulatory elements. **E, F.** Representative examples of CHiCAGO-and ABCC-detected contacts (for *SLU7* and *ZMAT2* promoters). The dashed line shows expected counts estimated using the CHiCAGO distance function. PIRs detected with CHiCAGO at 5 kb resolution are shown as red dots and shading, with ABCC as blue dots and shading and by both approaches as green points and shading. Black filled dots represent imputed counts considered by ABCC, corresponding to the maximum value between observed and expected counts. Unfilled dots represent observed counts falling below expected values.

Applying the ABCC algorithm to ILC3 PCHi-C data produced 18,877 putative enhancer-promoter pairs across 17,690 genes (**Fig. S2F; Data S4** at https://osf.io/aq9fb). Similarly to CHiCAGO-detected PIRs, there was a positive association between the number of ABCC enhancers and gene expression (**Fig. 2B**). However, ABCC-detected interactions generally spanned shorter distances than CHiCAGO-detected pairs (median distance ∼69 kb vs ∼108 kb, respectively, p-value < 2.2e-16, Wilcoxon rank-sum test) (**Fig. 2C**), and the two sets of contacts showed only a limited overlap (8.4%; **Data S5** at https://osf.io/aq9fb). Nonetheless, as expected, both CHiCAGO PIRs and ABCC enhancers were enriched for active and open chromatin features, as well as CTCF binding sites and/or annotated motifs (**Fig. 2D**). Representative examples of jointly detected regulatory landscapes are shown in **Fig. 2E**. We combined ABCC- and CHiCAGO-detected promoter contacts for downstream analyses, referring to them collectively as PIRs hereafter.

### Comparative analysis of promoter interactomes between ILC3 and CD4+ T cells identifies shared and differential regulatory circuitries

ILC3s share developmental similarities^51,52^ and common “immune modules” with CD4+ T cells^52–54^, prompting us to use this abundant cell type for comparative analysis and identification of ILC3-specific regulatory circuits. To this end, we generated and processed high-resolution PCHi-C data for CD4+ T cells using the same protocol, identifying 31,252 and 87,348 interactions at single-fragment and 5 kb resolution, respectively (**Data S6** and **S7** at https://osf.io/aq9fb). In addition, we detected 30,258 enhancer-gene pairs with ABCC across 16,956 genes (**Data S8** and **S9** at https://osf.io/aq9fb), 30% of which were shared with ABCC pairs identified in ILC3s. Differential analysis of chromatin interactions between ILC3s and CD4+ T cells with Chicdiff^55^ revealed a total of 19,038 cell-type-specific interactions (1,818 at fragment resolution and 17,220 at 5 kb resolution) across 3,664 genes (weighted adjusted p-value <0.05) (**Fig. 3A**). As expected, we also detected a significant association between differential interactions and differential expression (chi-squared = 23.938, df = 1, p-value = 9.948 x 10^-7^) (**Fig. 3B**; **Data S10** at https://osf.io/aq9fb).

**Figure 3.**
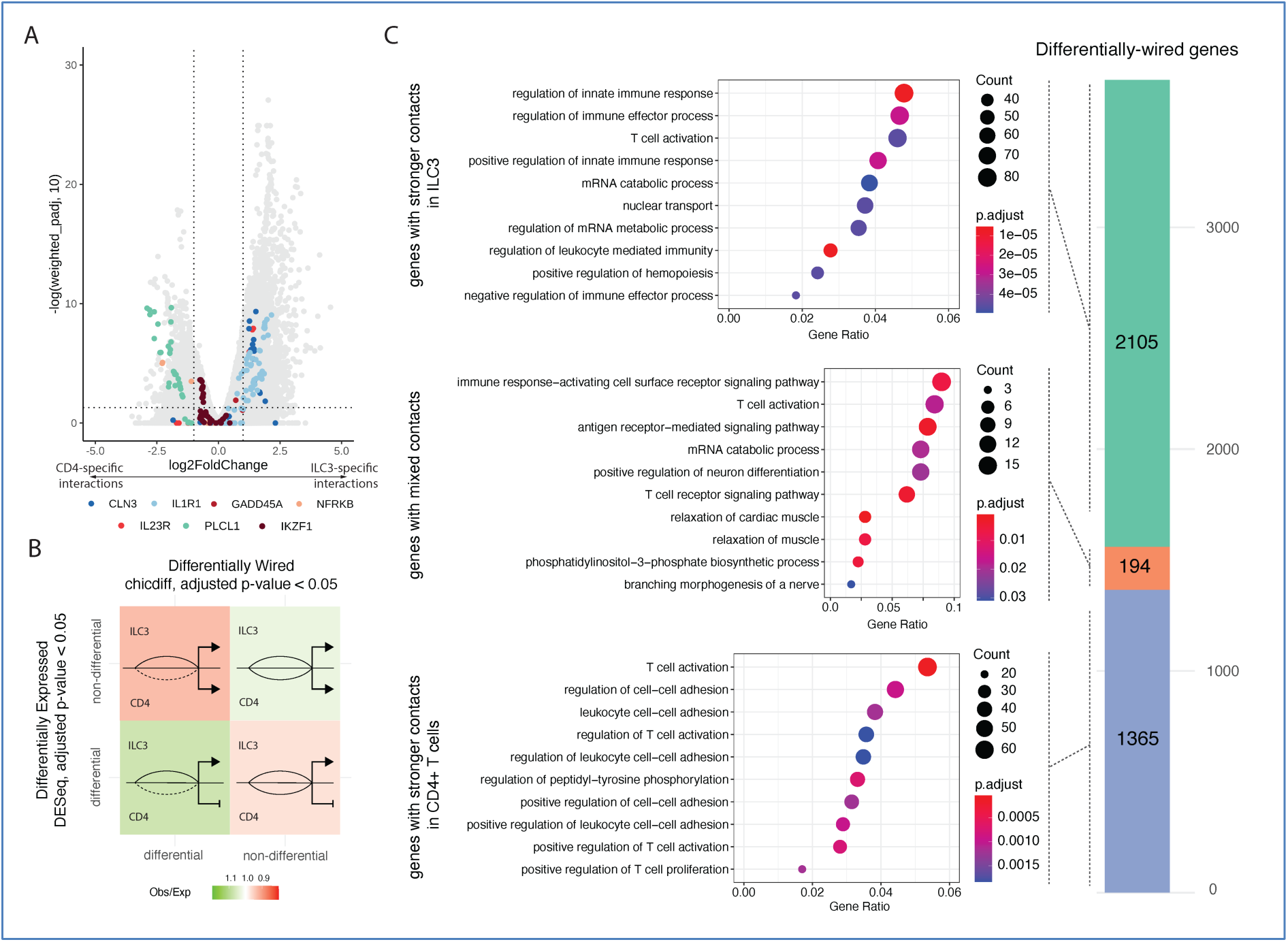
Differential enhancer-promoter interactions between ILC3s and CD4+ T cells. **A.** Volcano plot of differential interactions between ILC3s and CD4+ T cells detected by Chicdiff, highlighting those of selected immune-related genes (*CLN3*, *IL1R1*, *GADD45A*, *NFKB*, *IL23R*, *PLCL1*, *IKZF1*). **B.** Relationship between differential expression (DESeq2, adjusted p < 0.05) and differential wiring of promoter contacts (Chicdiff, adjusted p < 0.05). **C.** Gene Ontology enrichment analysis of genes with stronger contacts in ILC3s (top), CD4+ T cells (bottom) or a mixture of contacts that are stronger in either cell type (middle), showing biological processes related to immune cell activation, adhesion, and differentiation. Bubble size reflects the number of genes; colour indicates adjusted p-values. The bar plot shows the overlap between differentially wired genes (as evaluated by Chicdiff) in ILC3s and CD4+ T cells.

Genes with increased ILC3-specific chromatin contacts were enriched for annotation terms such as “regulation of innate immune response,” including *NFKB1* (NF-κB signaling), *TLR3* (innate immune receptor), and *IFNG* (effector cytokine), and “regulation of immune effector process”, including *IL23R* (controlling ILC3 activation and cytokine production), *IL1R1*, *TNFSF4*, and *SOCS5* (negative feedback on cytokine signalling) (**Fig. 3C; Fig. S3A; Table S2**). In contrast, genes with CD4+ T cell-specific contacts were involved in “regulation of T cell activation” (e.g. *CD3E, CD86, CTLA4, IL6, FOXN1*) and “negative regulation of the MAPK cascade” (e.g. *DUSP14, DUSP16, PTPN6*) (**Fig. 3C; Fig. S3B; Table S3**).

We also identified 194 genes with differential contacts between ILC3s and CD4+ T cells, including *BCL2, FYN, CD226* (activating receptor on T and NK/ILC3-like cells), and *CCR7* (guiding ILC3 positioning and migration) (**Fig. 3C; Fig. S3C; Table S4**). Notably, many genes with ILC3- and/or CD4+ T cell-specific contacts converged on pathways such as TCR signalling and T cell activation (e.g. *IL23R, RORC, NFKB1, CD300A, PIK3R1, ZAP70, CTLA4, CD3E, CD226, ITK, CD28, CCR7*), indicating differences in the regulatory wiring of these genes in ILC3s and their adaptive immune counterparts. In contrast, genes with similar contact profiles across both cell types were associated with processes such as histone modification, chromatin remodelling, and lymphocyte proliferation and differentiation (**Fig. S3; Table S5**), reflecting their shared functionality in both cell types.

In conclusion, our comparative chromosomal interaction analysis highlights both shared and distinct regulatory wiring of ILC3s and CD4+ T cells, reflecting their specialised roles in innate versus adaptive immune responses and coordinated regulation of immune activation pathways.

### Promoter-interacting regions in ILC3s and CD4+ T cells are enriched for genetic variants associated with autoimmune disorders

Genetic risk variants for complex diseases are strongly enriched at transcriptional enhancers^14–16^. Therefore, we investigated whether regulatory elements interacting with gene promoters in ILC3s and CD4+ T cells were enriched for genetic susceptibility to human traits and diseases, using the RELI algorithm^56^ (**Fig. 4A**; see Methods). Briefly, RELI determines significantly enriched overlaps between selected genomic loci (here, promoter-interacting regions intersecting open chromatin or H3K27ac signals in ILC3s based on public data) and trait-associated genetic variants. This is done by comparing the observed overlaps with a null distribution of artificially created variant sets with similar linkage disequilibrium (LD) characteristics to the trait-associated variants^56^. A practical advantage of RELI over the commonly used stratified LD score regression^57^ is that it does not require summary statistics data and can be performed on sets of significant SNPs reported in the GWAS Catalog^58^.

**Figure 4.**
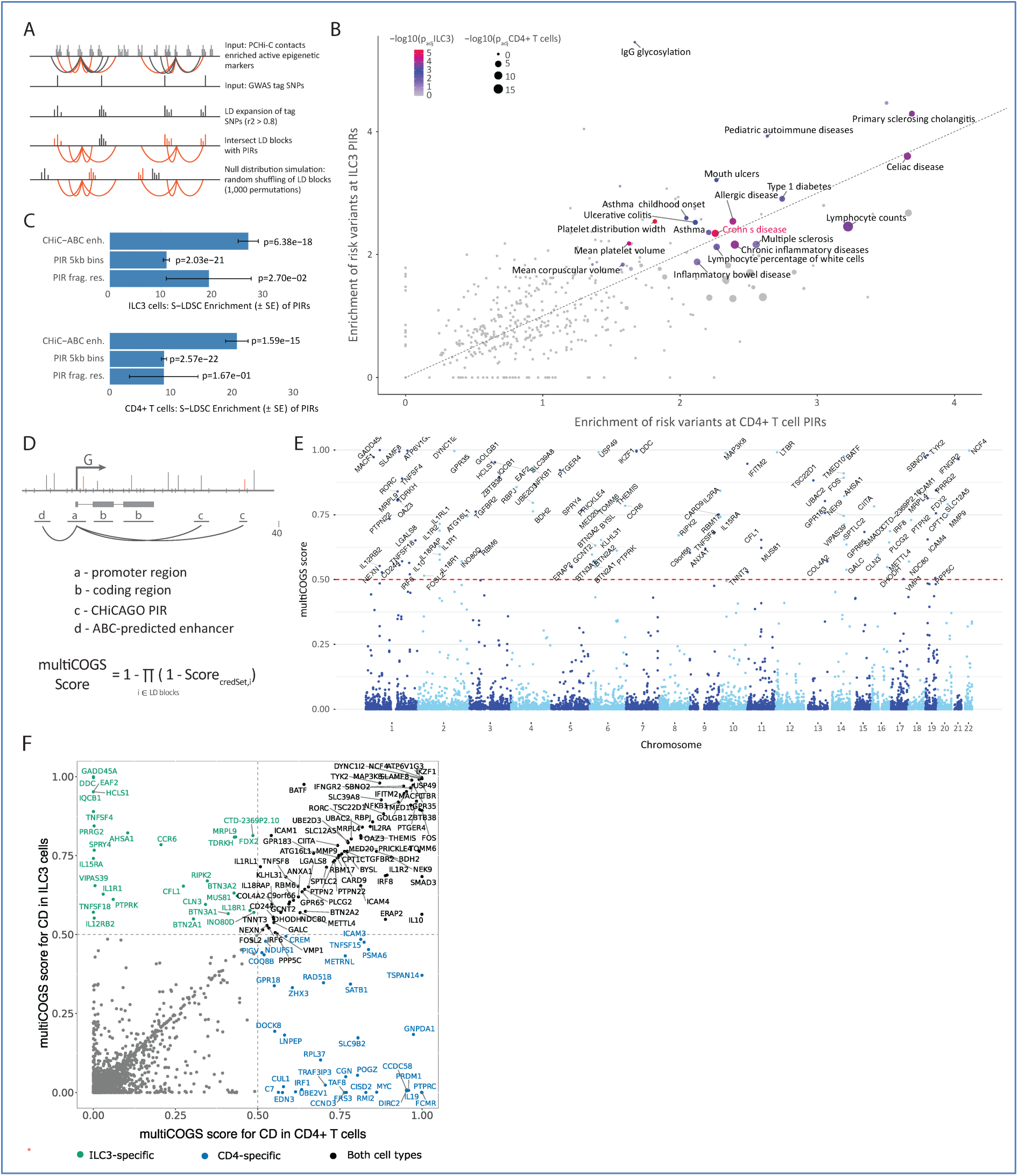
Statistical integration of PCHi-C results in ILC3s and CD4+ T cells with GWAS enables gene prioritisation for Crohn’s disease (CD) **A.** Schematic of the RELI algorithm used for estimating the enrichment of genetic risk loci within PIRs. **B.** RELI enrichment of risk variants in ILC3 vs CD4+ T cell PIRS across 495 diseases and traits. Traits with log_10_(BH corrected p-value in ILC3s) < 0.05 are labelled. **C.** Stratified LD score regression analysis for enrichment of CD risk heritability at PIRs of ILC3s and CD4+ T cells. **D.** Schematic of the multiCOGS algorithm. **E.** Manhattan plot of multiCOGS gene prioritisation scores for CD risk based on GWAS integration with promoter interactions in ILC3s. Genes with multiCOGS scores above 0.5 are labelled. **F.** Comparison of multiCOGS scores for CD obtained with promoter interactions detected in ILC3s and CD4+ T cells. Prioritised genes are labelled in green (multiCOGS scores > 0.5 in ILC3s only), blue (multiCOGS scores > 0.5 in CD4+ T cells only) and black (multiCOGS scores > 0.5 in both cell types). All other genes are shown as grey dots.

Out of the 495 analysed traits and diseases tested from the GWAS Catalog, genetic risk loci for 21 human traits were significantly enriched at promoter-linked putative regulatory elements in ILC3s (BH adjusted p-value < 0.05; **Fig. S4A**, **Table S6**; see Methods). Autoimmune diseases were overrepresented among these traits (according to the ontology EFO:0005140; p-value = 1.077 x 10^-5^, hypergeometric one-tailed test), affecting a broad array of organs and tissues that ILC3s are known to reside in. These included the gut (CD, celiac disease, ulcerative colitis, primary sclerosing cholangitis), airways (asthma, hay fever), and the central nervous system (multiple sclerosis). We also noted several traits of peripheral blood cells, including platelet width, lymphocyte count, and corpuscular volume (**Table S6**).

In CD4+ T cells, 22 traits were significantly enriched at promoter-interacting regulatory elements of CD4+ T cells (BH adjusted p-value < 0.05; **Fig. S4B**), with significant correlation between the two cell types (R^2^ = 0.845822, df = 10, 95% CI (0.5284, 0.9558), p = 0.00052; **Fig. 4B**), in line with the assumption that CD4+ T cells and ILC3 cells share many cis-regulatory circuits. However, several traits displayed cell-type specificity, such as allergic sensitisation, mouth ulcers, and IgG glycosylation in ILC3s, and primary biliary cirrhosis, rheumatoid arthritis, and systemic lupus erythematosus in CD4+ T cells (**Table S6**).

Among the autoimmune disorders, CD risk variants were particularly highly enriched within the active PIRs of both ILC3s and CD4+ T cells (∼2.3-fold enrichment in both cell types, p-value = 1.41 x 10^-8^ in ILC3s and p-value = 2.41 x 10^-10^ in CD4+ T cells). We confirmed this observation using stratified LD score regression (**Fig. 4C**). While the critical role of CD4+ T cells in CD is well-established^59–62^, the connection between ILC3s and CD pathogenesis is more recent. ILC3s are thought to influence inflammatory processes in CD, such as GM-CSF signalling and overexpression of the cytokines IL-22, IL-17, and IFN-γ^11,63^. We next sought to leverage PCHi-C data to prioritise genes linked to CD risk variants in these cell types.

### MultiCOGS prioritises genes linked to Crohn’s disease risk based on multivariate fine-mapping of imputed GWAS signals and promoter contacts in ILC3 and CD4+ T cells

To identify putative causal variants and genes for CD in ILC3s and CD4+ T cells, we extended our previously published Bayesian prioritisation algorithm, COGS^30,31^, which provides a single measure of support (“COGS score”) for each gene’s association with a trait of interest, calculated based on the location of fine-mapped GWAS signals within (i) gene coding regions, (ii) gene promoters, and (iii) promoter-interacting regions.

Despite its demonstrated utility in prioritising gene candidates in a range of human traits^30,31,64,65^, we identified areas for improvement in COGS. First, if the summary statistics underlying the trait-associated loci are too sparse, COGS may miss likely causal variants intersecting promoter-interacting regions. To mitigate this, we imputed additional trait-associated variants using an established summary statistics-based methodology^66^. Second, the original statistical fine-mapping approach utilised in COGS assumes at most a single causal variant per linkage disequilibrium (LD) block, whereas the latest evidence suggests that trait-associated LD blocks can contain multiple causal variants^67^. To address this, we updated the COGS algorithm to enable integration with recently developed multivariate fine-mapping approaches, such as SuSiE^68–70^ (**Fig. 4D**; see Methods). Finally, we accounted for both CHiCAGO- and ABCC-detected promoter-interacting regions. We refer to the updated version of COGS as “multiCOGS”.

We ran multiCOGS on the CD GWAS meta-analysis by de Lange *et al.*^71^ using the compendium of CHiCAGO- and ABCC-detected promoter-interacting regions in ILC3s or CD4+ T cells. At the previously established COGS score cutoff of 0.5^30^, we prioritised 109 genes in ILC3s (**Fig. 4E**) and 118 genes in CD4+ T cells (**Fig. S5A**; **Table S7**). The majority of genes were prioritised based on 3D proximity of non-coding trait-associated variants to gene promoters, either by PCHi-C or ABCC (**Fig. S5B**). ABCC contributed to around 11% of the prioritised genes in both cell types (**Fig. S5C**). At first examination, we noted many candidate genes with roles in immune processes already known to be dysregulated in inflammatory bowel disease (IBD)^72–74^. Examples include cytokine signalling (*IL10*, *IL1RL1, LTBR, IL2RA, IFNGR2, TNFSF8*), autophagy (*ATG16L1, GPR65*), and antimicrobial processes in the gut (*PTPN2, IRF8*)^75,76^. The prioritised genes also highlighted IL-23/Th17 signalling (for example, *RORC, NFKB1, IL2RA,* and *TYK2*), a known immune axis in CD pathology^77^, and known transcriptional regulators (*FOS*, *TSC22D1*, *RBPJ*). In several loci, multiCOGS prioritised several compelling gene candidates, based on multiple credible sets. For example, in ILC3s, two credible sets of variants in chr7p implicated the *IKZF1* gene (encoding the Ikaros transcription factor) by PCHi-C interactions, and the *DDC* gene (encoding dopamine regulator L-dopa decarboxylase) by ABCC pairing (**Fig. S5D**). Ikaros, an established critical regulator of immune cell development^78^, also scored highly in the original COGS algorithm. However, the more distal *DDC* gene, which has recently emerged as a potential regulator of immune cell infiltration^79^, scored well below the prioritisation threshold (**Table S7**). This demonstrates the potential of multiCOGS and ABCC for highlighting previously missed gene candidates.^79^

We next explored more closely how the results of multiCOGS compared with those from our previously published COGS pipeline, which used univariate fine mapping without imputation and was based purely on CHiCAGO results without ABCC (hereafter referred to as “classic COGS”). Classic COGS resulted in substantially smaller prioritised gene sets (55 genes in ILC3 cells and 75 genes in CD4+ T cells with COGS score > 0.5) (**Table S7**). As examples, we note that compelling candidate genes such as *IL12RB2* and *IL15RA* (in ILC3s), *TNFSF15,* and *ICAM3* (in CD4s), and *NFKB1*, *BATF*, *ICAM1* and *TNFSF8* (in both cell types) were only prioritised in multiCOGS (**Table S7**). Moreover, we discovered that both of the novel aspects of multiCOGS (imputation and multivariate fine mapping) contributed substantially to the increased number of genes prioritised in comparison with classic COGS (**Fig. S6A**). For the majority of genes, multiCOGS prioritisation scores were similar or higher than in conventional COGS in both ILC3s and CD4s (**Fig. S6B**). Only five genes prioritised by conventional COGS had sub-threshold scores in multiCOGS, including *JAK2* (see **Fig. S6C** and **Supplementary Note 1**).

Next, we searched for prior evidence of association of all multiCOGS-prioritised genes with CD (or IBD, more broadly) by querying the top CD genes in OpenTargets, curated gene-to-disease databases, and functional studies^37,80–83^. We found that over half of multiCOGS-prioritised genes in ILC3s (61/109) and CD4s (67/118) were not previously implicated in these databases (**Table S8**). These newly prioritised genes included compelling candidates such as ubiquitin-specific peptidase 49 (*USP49*), adding to the existing evidence for the role of protein ubiquitination in IBD development^84^, and lymphotoxin beta receptor (*LTBR*), known to be important for gut epithelial cell IL-23 production^85^. In particular, 23 genes selectively prioritised in ILC3s **(Fig. 4F)** were not previously linked to CD in the studied datasets. These included genes with unexpected functions, such as the neurotransmitter DOPA decarboxylase (*DDC*), and a lysosomal/endosomal transmembrane protein (*CLN3*). *CLN3* is involved in lipid trafficking and catabolism^86,87^, and mutations in this gene cause Batten disease, a group of lysosomal storage disorders characterised by progressive neurodegeneration^88^.

Taken together, by accounting for imputed variants and multiple causal variants per locus, multiCOGS expands the ability to discover candidate genes in complex trait loci using promoter interactions.

### Prioritised gene candidates in ILC3 cells implicate inflammatory processes in CD aetiology

We explored the biological functions of the 109 prioritised CD genes in ILC3s based on their public gene set annotations (**Table S9**). Seven biological states or processes were significantly enriched among the gene candidates: IL6-JAK-STAT3 signalling, TNFα signalling via NFκB, IL2-STAT5 signalling, inflammatory response, allograft rejection, IFNγβ response, and TGFβ signalling (Hallmark gene sets; **Fig. S7A**). Molecular functions included cytokine receptor activity and NAD+ metabolic activity (GO Term Molecular Functions, **Fig. S7B).** We saw the strongest enrichment of cell-type signatures for tissue-resident immune cells, including gastric and duodenal immune cells, as well as monocytes, dendritic cells, and basophils in the lung (**Fig. S7C**). We also noted the signature for ILC progenitor cells in fetal lung^89^, driven by the genes *IL1R1, ICAM1, IFNGR2, PLCG2, CCR6,* and *RORC* (adjusted p = 0.0176). Enriched curated pathways highlighted immune-mediated diseases, including rheumatoid arthritis, neuroinflammation, IBD, and bacterial infection (WikiPathways; **Fig. S7D**). Other relevant pathways included T cell differentiation and signalling of IL-18, a key cytokine for ILC3 function^90^ (**Fig. S7C**). Leveraging published IBD patient gene sets^91^, we also found enrichment for genes differentially expressed in the rectum in patients with CD (adjusted p-value = 1.38 x 10^-4^) and ulcerative colitis (adjusted p-value = 0.0156) (**Table S9**, **Fig. S7E**).

We then investigated which transcription factors (TFs) might regulate the CD gene candidates in ILC3s using two methods. First, we used a gene-centric approach to identify overrepresented genes predicted to be targeted by a given TF (TF targets from MSigDB). This analysis highlighted the architectural protein HMGA1 and the known inflammatory response regulator NFκB (**Table S9** and **Fig. 5A**). Second, we used a region-centric approach, searching for enrichment of predicted TF binding sites across a range of cell types at the PIRs of CD candidate genes in ILC3s. We found significant enrichment for 97 TFs (**Fig. 5B, Table S10**), many of which were previously implicated in inflammatory response, including IKZF1/Ikaros^92^, BATF^93^, and NFKB3/RELA^94–96^, which are all highly expressed in ILC3s **(Fig. 5C)** and have established roles in ILC3 biology. Two examples of potential long-range regulation of CD candidate genes by putative TF binding at PIRs are shown in **Fig. 5D** and **E**. In the first example, the promoter of the *IKZF1* gene contacts two upstream PIRs, each containing a separate credible set of fine-mapped CD susceptibility variants and bearing marks of open and active chromatin (ATAC-seq and H3K27ac peaks) in ILC3s. Based on data from lymphoblastoid cell lines, these PIRs recruit multiple TFs: IKZF1 itself, as well as BATF, NFKB3, ATF2, and the architectural proteins CTCF and SA1 (**Fig. 5D**). In the second example, the promoter of *IL1R1* contacts CD risk variant-containing PIRs that have accessible chromatin in ILC3s and contain CTCF binding signals in lymphoblastoid cell lines (**Fig. 5E**).

**Figure 5.**
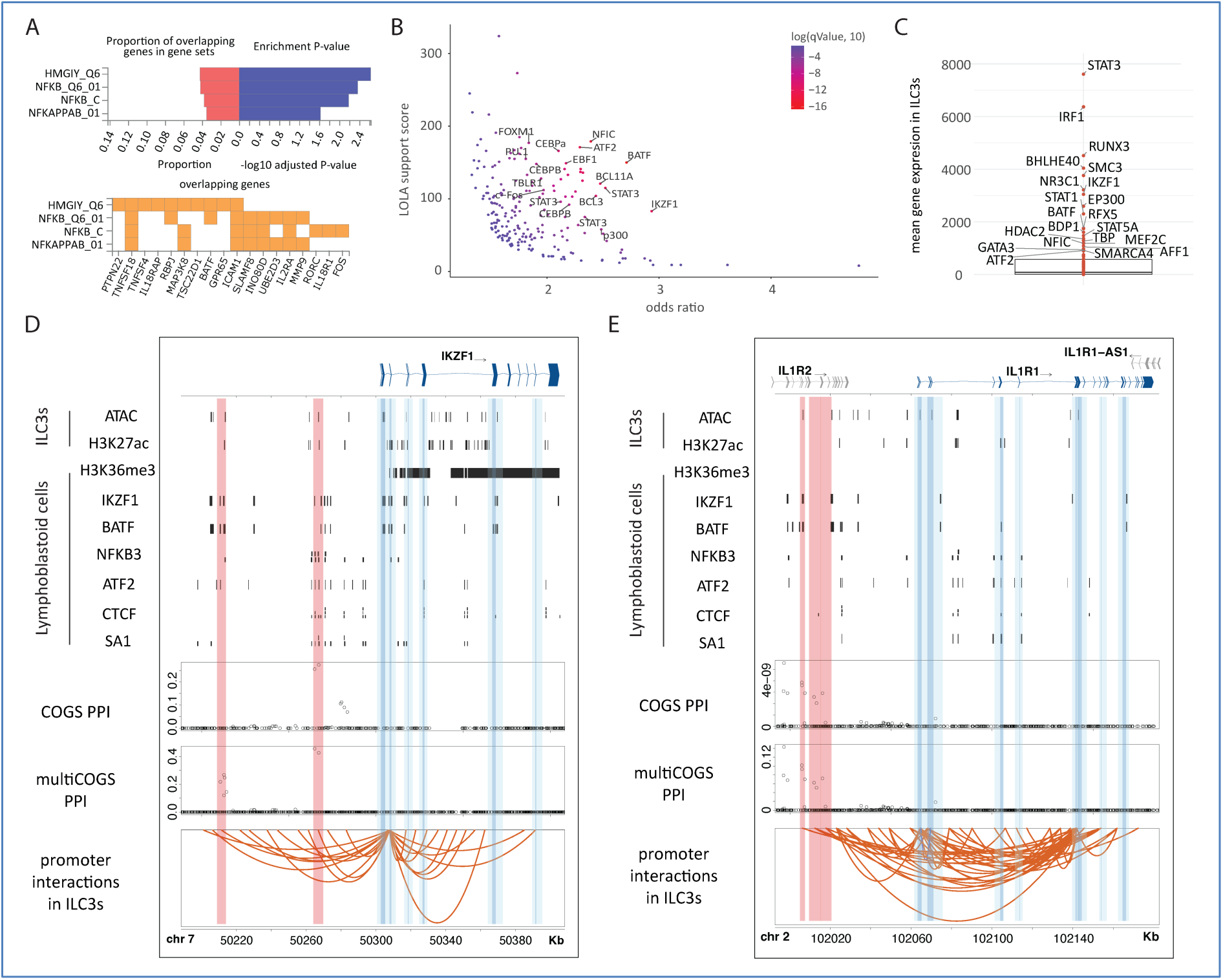
Characterisation of genes associated with CD risk prioritised by multiCOGS in ILC3s and their putative TF regulators. **A.** Significant sets of multiCOGS-prioritised genes predicted to bind specific TFs in their promoter regions, according to the MSigDB TF targets database, detected using the GENE2FUNC pipeline in FUMA^121^. TF sets are labelled (rows), with the proportion of all multiCOGS genes per set and the associated p-values shown on the top panel, and the gene names on the bottom panel. **B.** Enrichment analysis for TF binding sites at active PIRs for genes prioritised by multiCOGS vs active PIRs of all genes submitted to multiCOGS analysis. **C.** Expression of TFs enriched at the PIRs of prioritised genes. Outliers are removed for clarity. **D and E.** Examples of genes prioritised by multiCOGS for CD (*IKZF1*, and *IL1R1*), showing patterns of TF binding in lymphoblastoid cell lines, and posterior probability profiles of classic COGS and multiCOGS. Vertical dark blue and light blue bands, respectively, highlight annotated gene promoters and promoter-proximal regions (+/− 5 restriction fragments) considered in (multi)COGS analysis in addition to PIRs. Vertical red bands highlight PIRs harbouring CD risk-associated SNPs with high posterior probability of inclusion. Orange arcs correspond to significant interactions (CHiCAGO score > 5) at 5kb resolution for *IKZF1* (E) and *IL1R1* (F), respectively.

Jointly, these results propose inflammatory signalling genes as causal candidates for CD susceptibility in ILC3s.

### *CLN3* contributes to ILC3 inflammatory capacity

We next focused on *CLN3,* a gene implicated in the neurodevelopmental disorder Batten disease. *CLN3* was selectively prioritised as a CD risk gene in ILC3s, but not CD4+ T cells, and has not previously been linked to CD or other immune-mediated diseases. Examination of the SuSIE fine-mapped CD GWAS locus underlying *CLN3*’s prioritisation revealed a credible set of variants overlapping two regions considered by multiCOGS. The first region is an ILC3-specific *CLN3* PIR located 14.2 kb downstream of the canonical *CLN3* TSS (red band in **Fig. 6A**). The second region lies between exons 10 and 11 of the canonical *CLN3* transcript, adjacent to an annotated internal promoter (first dark blue band in **Fig. 6A**). Unexpectedly, we found that both regions lacked chromatin accessibility and enhancer activity signals in ILC3s, as well as in all other cell types included in the Ensembl Regulatory Build database (**Fig. 6A**). Data from lymphoblastoid cell lines^97^ showed enrichment for the H3K36me3 mark, which is typically associated with transcriptional elongation^98^ and facultative heterochromatin^99^ (**Fig. 6A**). To seek complementary evidence for a regulatory role of this locus, we queried the OpenTargets database^100^ for possible colocalisation between the CD risk signal and known *CLN3* expression quantitative trait loci (eQTLs). CD risk GWAS and *CLN3* expression were likely to share a joint causal genetic signal (posterior probability ≥ 0.8, as determined by coloc^101^ and reported in OpenTargets) in whole blood^102,103^, monocytes^104,105^, thyroid^103^, small intestine^103^, and cerebellum^103^. Notably, the same CD GWAS signals also colocalised with eQTLs for nearby genes, including *APOBR*, which is located ∼2 kb downstream of *CLN3* in a divergent orientation, suggesting a complex regulatory architecture at this locus.

**Figure 6.**
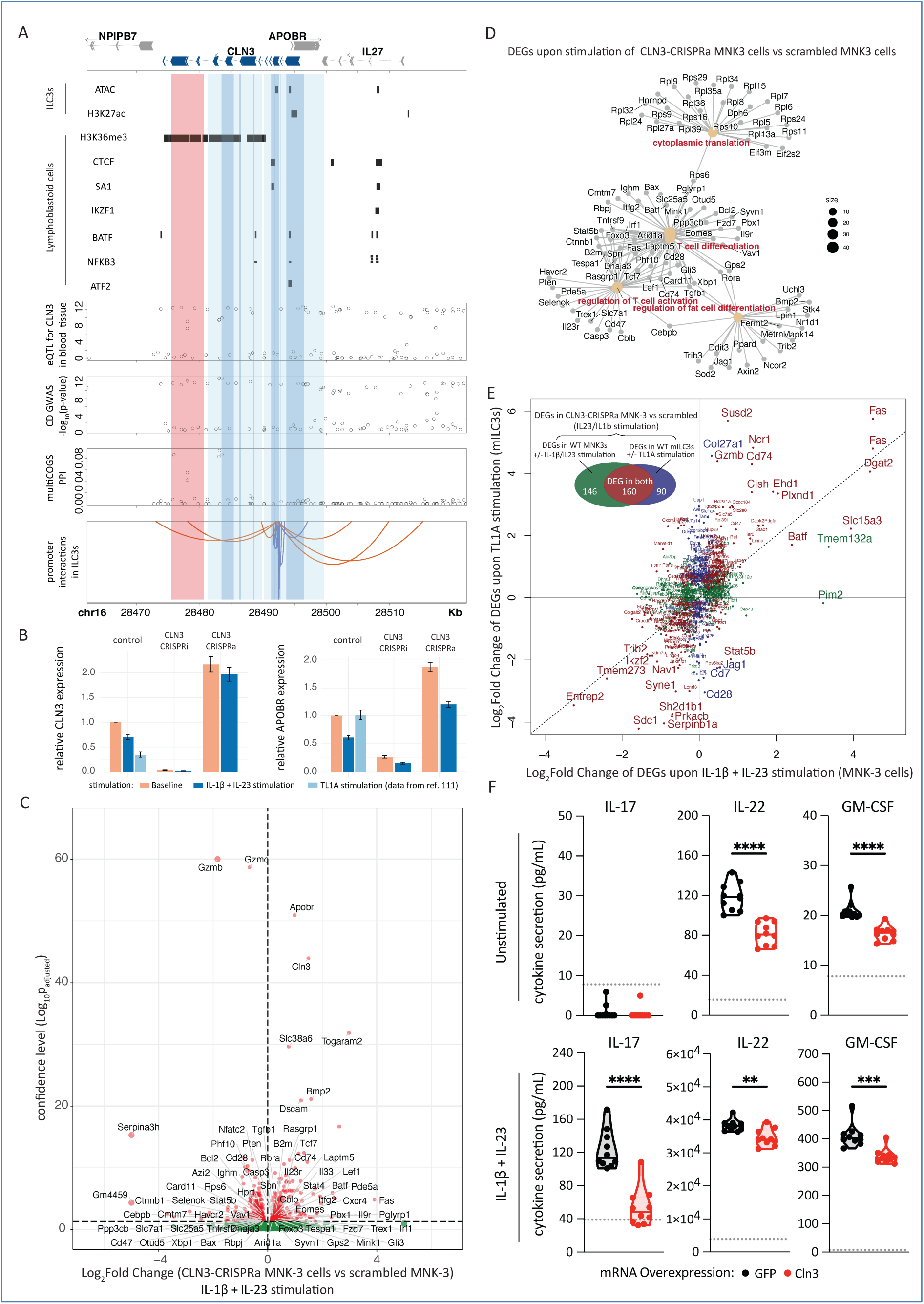
Evidence for the role of CLN3 in ILC3 inflammatory function. **A.** Interaction profile of the human *CLN3* promoter alongside the tracks of TF binding, blood eQTLs, CD GWAS and SuSiE posterior probabilities of inclusion. Dark blue and light blue bands, respectively, highlight the locations of annotated *CLN3* promoters and promoter-proximal regions (+/− 5 restriction fragments) considered by multiCOGS in addition to PIRs. Red band highlights the ILC3-specific PIR containing CD-associated SNPs with high posterior probability of inclusion. Orange and purple arcs, respectively, depict significant interactions (CHiCAGO score > 5) in ILC3s at 5kb and single-fragment resolution. **B.** Up- and down-regulation of *Cln3* and *Apobr* upon TL1A stimulation in mouse primary ILC3s (RNA-seq data from Ref.^108^) and upon IL-23/IL-1β stimulation in CLN3-targeted CRISPRi and CRISPRa MNK-3 cells (RNA-seq data from this study). **C.** Differential expression of genes in IL-23/IL-1β-stimulated *Cln3*-CRISPRa MNK-3 cells relative to scrambled gRNA controls. Red dots - differentially expressed genes (stimulated *Cln3-*CRISPRa DEGs, DESeq2 adjusted p-value < 0.05), with other genes shown as green dots. **D.** Network-style representation of GO term enrichment analysis of stimulated *Cln3-*CRISPRa DEGs. **E.** Changes in the expression of stimulated *Cln3-*CRISPRa DEGs (dots) upon either IL-23/IL-1β or TL1A stimulation of unperturbed MNK-3 cells (data from Ref.^108^). **F.** Evidence that *Cln3* overexpression decreases inflammatory cytokine secretion. MNK-3 cells were electroporated with GFP mRNA (black) or Cln3-myc mRNA (red), then cultured either unstimulated (top row) or stimulated with IL-1β and IL-23 (bottom row) for 24 hr. Cytokine concentrations (IL-17, IL-22, GM-CSF) in culture supernatants were quantified by ELISA. Each point represents an individual biological replicate (n=10 per condition). The data shown are from one representative experiment of three independent experiments performed. Dotted line indicates the lower limit of quantification for each assay. Statistical significance was assessed using an unpaired Welch’s t-test. p<0.01 (**), p<0.001 (***), p<0.0001 (****).

To further investigate the role of the *CLN3* locus in ILC3s, we used mouse MNK-3 cells as a tractable model for ILC3 activation and effector function. We found that *Cln3* expression was downregulated upon stimulation of MNK-3 cells with IL-23 and IL-1β, cytokines that are essential for ILC3 effector function^106,107^ (**Fig. 6B**, left). Consistent with this observation, analysis of published RNA-seq data from primary mouse ILC3s stimulated with TL1A^108^ also showed reduced *Cln3* expression (**Fig. 6B**, left). Notably, the adjacent gene *Apobr* was similarly downregulated under IL-23/IL-1β stimulation (**Fig. 6B**, right), in line with eQTL-based evidence of coordinated regulation of these genes in humans.^102,103,104,105^ In contrast, TL1A stimulation did not affect *Apobr* expression (**Fig. 6B**, right).

To interrogate the transcriptional consequences of stimulation-induced *Cln3* repression, we used CRISPR activation (CRISPRa; dCas9-VP64 + MS2-p65-HSF1) to prevent *Cln3* downregulation in MNK-3 cells during stimulation. CRISPRa targeting produced an approximately threefold increase in *Cln3* expression in stimulated MNK-3 cells (**Fig. 6B**, left). Notably, *Apobr* expression was also increased in both basal and stimulated conditions (**Fig. 6B**, right), potentially reflecting local effects of CRISPRa targeting, but also mirroring the coordinated regulation observed at this locus (**Fig. 6B**). Bulk RNA-seq analysis revealed widespread transcriptional changes following *Cln3* CRISPRa, with 519 differentially expressed genes in unstimulated cells and 722 in stimulated cells relative to scrambled gRNA controls (DESeq2 adjusted p-value < 0.05; **Fig. 6C** and **S8A; Table S11** and **Data S10** at https://osf.io/aq9fb). These genes were enriched for pathways involved in lymphocyte differentiation, activation, and proliferation, including upregulation of *Cd23r*, *Cd74*, and *Fas*, and downregulation of the inflammatory serine proteases *Gzmb* and *Gzmc* (**Fig. 6D, Fig. S8B**). Notably, more than half of the genes differentially expressed in stimulated *Cln3-*CRISPRa cells overlapped with genes altered by IL-23/IL-1β or TL1A stimulation in *Cln3*-unperturbed cells^108^ (**Fig. 6E**), suggesting that sustained *Cln3* expression counteracts canonical activation-associated transcriptional programmes. In contrast, CRISPR interference (CRISPRi; dCas9-KRAB)-mediated knockdown of *Cln3* resulted in few transcriptional changes beyond *Cln3* and *Apobr* themselves **(Fig. S8C, D; Table S11; Data S11** at https://osf.io/aq9fb). Notably, these included upregulation of *Nos2,* a gene previously implicated in limiting ILC3-driven intestinal inflammation^107^.

Given the coordinated regulation of *Cln3* and *Apobr* expression upon ILC3 stimulation, the limited transcriptional impact of further *Cln3* knockdown in activated cells, and the pronounced effects of *Cln3* overexpression, we next asked whether the CLN3 protein modulates ILC3 effector function at a post-transcriptional level. CLN3 is a lysosomal and endosomal protein with established roles in vesicular trafficking, lysosomal homeostasis, and protein turnover^109,110,111^, processes that are central to cytokine storage and secretion. Therefore, we ectopically overexpressed *Cln3* in MNK-3 cells and measured cytokine secretion under basal and inflammatory conditions. Overexpression of the myc-tagged CLN3 construct was confirmed by RT-qPCR and immunoblotting (**Fig. S8E, F**). As expected, MNK-3 cells constitutively secreted IL-22 and GM-CSF, with further induction of these cytokines upon stimulation, whereas IL-17 production was restricted to stimulated conditions (**Fig. 6F** and **S8G**). Notably, CLN3 overexpression significantly reduced the secretion of IL-17, IL-22, and GM-CSF by stimulated MNK-3 cells (**Fig. 6F** and **S8G**). Basal IL-22 and GM-CSF secretion were also reduced in the absence of stimulation (**Fig. 6F** and **S8G**). Viable cell numbers were quantified at the end of cytokine secretion assays and showed no difference under basal conditions, with a modest reduction in Cln3-overexpressing cells following stimulation (**Fig. S8H**).

Collectively, these results highlight the Batten disease gene *Cln3* and the broader *Cln3/Apobr* locus as regulators of ILC3 inflammatory output, revealing a previously unrecognised role for this locus in shaping ILC function.

### MultiCOGS prioritises candidate genes for six autoimmune diseases with potential roles in ILC3 inflammatory function

Building on the methodologies and data generated in this study, we extended multiCOGS analysis in ILC3s and CD4+ T cells to five other autoimmune GWAS datasets in addition to CD with available summary statistics that showed enrichment at ILC3 PIRs in the RELI analysis: adult-onset asthma, IBD, ulcerative colitis (UC), primary sclerosing cholangitis (PSC) and celiac disease. Across the six traits and two cell types, we detected a total of 332 prioritised disease candidate genes (multiCOGS score > 0.5), of which 251 were prioritised in ILC3 cells (**Fig. 7A**) and 266 in CD4+ T cells (**Table S12**). As expected from their shared aetiology, the three traits relating to inflammatory bowel disease (CD, UC, and IBD) clustered together with respect to gene-level multiCOGS scores, while asthma formed an outgroup (**Fig. 7A**).

**Figure 7.**
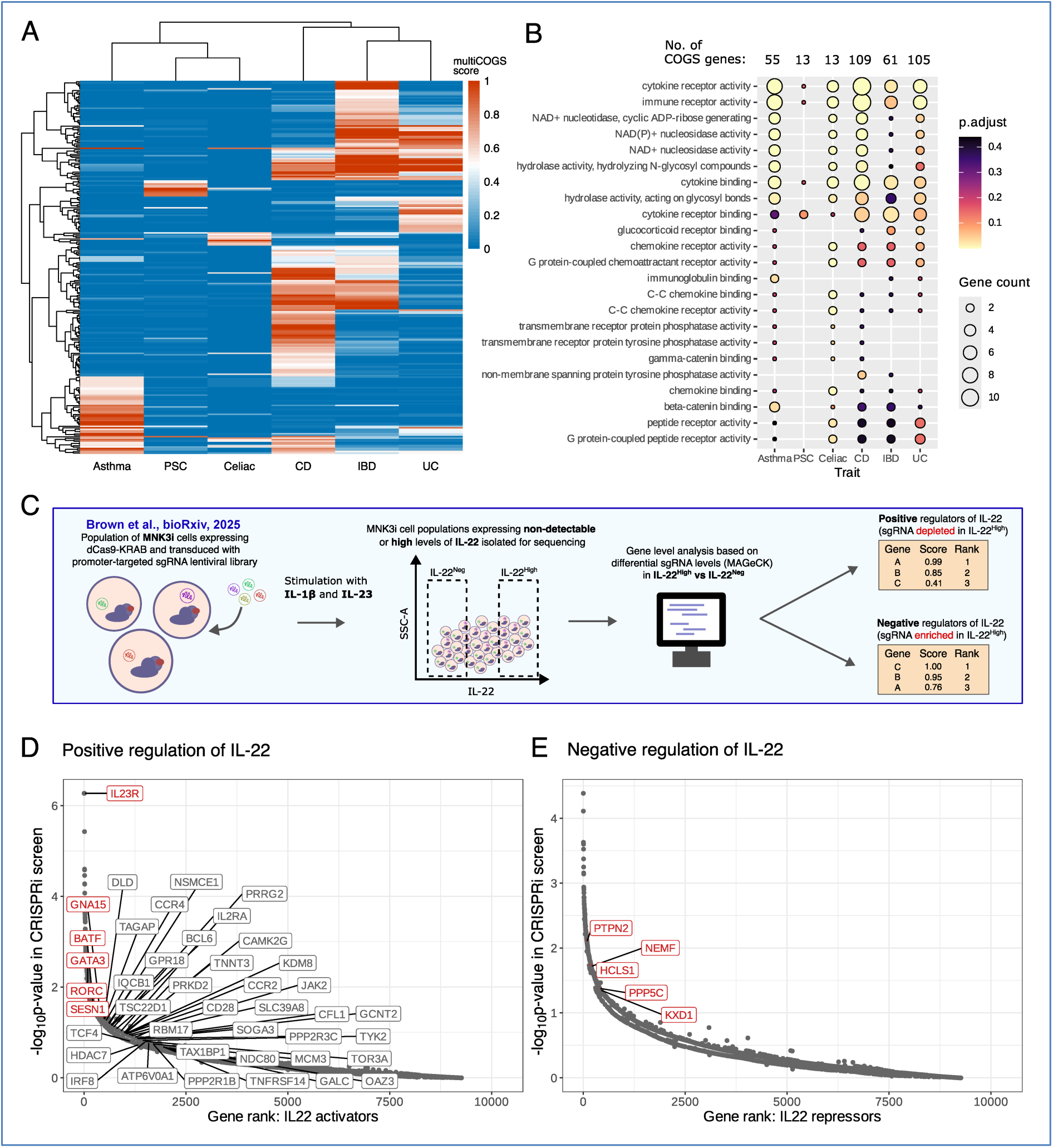
A compendium of prioritised genes in ILC3s for six autoimmune diseases. **A.** MultiCOGS results across asthma, primary sclerosing cholangitis (PSC), Celiac Disease, Crohn’s Disease (CD), Inflammatory Bowel Disease (IBD) and Ulcerative Colitis (UC) in ILC3 cells. Rows represent each gene that scored at least 0.5 in one of the traits. Colours show the multiCOGS score in each trait. Clustering on genes (rows) and traits per cell type (columns) is based on Euclidean distance. **B.** Significant hallmark pathways identified in at least one of the traits in ILC3 cells by GO term analysis. **C.** Schematic of the MNK-3 CRISPRi screen for detecting genes involved in the regulation of IL-22 signalling^125^. **D.** multiCOGS genes for all six traits visualised among the CRISPRi results, which are ranked by evidence of positive IL-22 regulation in the MNK-3i cells. The multiCOGS genes with p < 0.05 in the screen are labelled in red. MultiCOGS genes driving GSEA signal (“leading edge”) are labelled in grey. **E.** Similar to D, but for genes ranked by score for negative IL-22 regulation in the MNK-3i screen. Red labels indicate multiCOGS genes significant in the screen at p < 0.05. Since GSEA for multiCOGS genes among IL-22 repressors was not significant, the leading edge genes are not labelled. CD: Crohn’s Disease, IBD: Inflammatory Bowel Disease, GSEA: Gene Set Enrichment Analysis, PSC: primary sclerosing cholangitis, UC: Ulcerative Colitis.

A total of 66 candidate genes were prioritised in ILC3s only, and 81 in CD4+ T cells only (**Table S12)**. Notable ILC3-specific candidate genes included several cytokines and receptors involved in type I immune response, such as *CCR2* (celiac disease), *BCL6* and *IL17A* (both asthma), as well as the IL-18 receptor (*IL18R1*), which we previously prioritised for CD, and here also prioritised for celiac disease and asthma. We also noted family members of butyrophilin (BTN) proteins–immunomodulatory transmembrane proteins involved in recognition of microbial antigens–prioritised in both CD and asthma (*BTN3A1* and *BTN3A2*), specific to ILC3 cells. Finally, we noted that *CLN3* was prioritised for the broader IBD trait (multiCOGS score 0.538, **Table S12**) in addition to CD, again selectively in ILC3s.

Pathway analysis of the prioritised genes across the analysed traits revealed shared enriched GO terms for inflammatory processes such as cytokine binding and immune receptor activity (**Fig. 7B; Table S13A**). To gain further insight into the role of the prioritised genes in ILC3 inflammatory function, we turned to a recent CRISPRi screen for putative regulators of IL-22 expression in MNK-3 cells following IL-23/IL-1β stimulation^112^ (**Fig. 7C**). Of the multiCOGS gene candidates across all profiled autoimmune diseases, six were significant positive regulators and five were significant negative regulators of IL-22 protein production, as detected by the CRISPRi screen (**Table S13B** and labelled in red in **Fig. 7D** and **7E**). Among the IL-22 activators were three candidate genes for IBD-related traits, all with known strong roles in IL-22 activation (*IL23R*, *BATF,* and *RORC*). The remaining three IL-22 activators were all candidate genes for asthma alone: *GNA15*, *SESN1,* and *GATA3*, of which only *GATA3* has been previously reported to directly activate IL-22 in ILC3s^113^. Meanwhile, the five multiCOGS genes putatively downregulating IL-22 production were all associated with IBD-related traits (*PTPN2, NEMF, HCLS1, PPP5C, and KXD1*). Of these, only *PTPN2* has direct evidence for IL-22 repression, through STAT3 dephosphorylation^114^. The other putative IL-22 negative regulators have diverse functions in protein homeostasis (*NEMF*), actin remodelling (*HCLS1*), stress signalling (*PPP5C*), and lysosome localisation (*KXD1*). Overall, multiCOGS genes were significantly enriched among the genes scoring highly for positive IL-22 regulation (GSEA p = 0.0284, **Table S13C**; genes driving the association labelled in **Fig. 7D**), implicating the control of ILC3 activation as an important mechanism underpinning the effects of the prioritised genes on autoimmune disease risk.

In summary, this analysis expands the compendium of prioritised GWAS gene candidates with potential roles in ILC3s to six autoimmune disease traits and demonstrates the potential role of many prioritised genes in ILC3 inflammatory function.

## Discussion

In this study, we present high-resolution promoter interaction profiling in ILC3s, revealing tens of thousands of promoter contacts with enhancers and GWAS variants associated with multiple immune diseases, including those that are unique to ILC3s compared with their phenotypically related counterparts in the adaptive immune system, CD4+ T cells. ILC3s are a relatively rare cell type that cannot be easily expanded *in vivo*, which makes their chromosomal interaction profiling challenging. Indeed, this problem precluded ILC3 profiling by standard Hi-C alongside type 2 ILCs in a recent mouse study^115^. Robust Capture Hi-C profiling typically requires even higher cell numbers. Our efficient PCHi-C protocol^44^ and the use of a four-cutter enzyme (*DpnII)* have enabled a higher-resolution analysis of human ILC3s in this study, adding these clinically-relevant cells to the ever-expanding array of cell types with available promoter interactome maps, including the 17 abundant blood cell types that we profiled previously using high-coverage PCHi-C at a six-cutter enzyme (*HindIII)* resolution^30^. While emerging technologies provide complementary solutions for the inference of enhancer-promoter relationships, such as through the correlated activities of these elements across cell types or single cells, genetic evidence and high-throughput perturbation screens, 3D genomics-based approaches continue to offer unique advantages by delivering mechanistically-grounded information in high throughput at a reasonable cost and time investment.

Unlike in our previous studies, here we take advantage of two conceptually different computational analysis strategies for detecting promoter contacts from Capture Hi-C data. The first strategy is based on our established CHiCAGO pipeline to detect ‘significant contacts’ – i.e., those whose frequency significantly exceeds the expectation at a given distance and technical noise levels. The second strategy is based on the adaptation of the ABC approach^14,41^ to Capture Hi-C data (the Activity-by-Captured-Contact method, ABCC), which, in contrast, considers the raw contact frequency rather than its significance. As expected from this conceptual difference, ABCC prioritises shorter-range contacts compared with CHiCAGO, resulting in the largely non-overlapping sets of identified contacts and GWAS-prioritised genes. However, the longer-range contacts detected using CHiCAGO, which were also enriched for active enhancers, drive the majority of our identified disease associations. From the practical point of view, therefore, these two approaches are largely complementary, and their combined use is warranted. Mechanistically, this suggests that at short linear distances, the background frequencies of promoter-enhancer contacts arising from constrained Brownian motion are sufficient for the functional interactions between these regions. In contrast, at longer ranges, additional factors (e.g., cohesin-mediated loops) are likely required to facilitate the statistically unusual contact frequencies and enable functional interactions.

We find a strong enrichment for CD-associated SNPs within the ILC3 PIRs, consistent with recent findings showing that superenhancers specific to ILC3 or Th17 cells, rather than to ILC1 or Th1 cells, preferentially contain CD-associated variants^21^. Using our multiCOGS strategy that integrates GWAS data processed with multivariate statistical fine-mapping with information on enhancer-promoter links from PCHi-C, we prioritise a total of 109 genes in ILC3s, 29 of which are not detected in CD4+ T cells. Notably, the number of multiCOGS-prioritised genes has increased considerably compared with the results obtained with our previously developed COGS pipeline^30,31^. The key improvements of multiCOGS include summary statistics-based imputation and allowing for multiple causal variants per linkage disequilibrium (LD) block. At the molecular level, the increased recall of prioritised genes reflects the fact that the same LD block often contains multiple regulatory elements (including promoter-proximal and distal enhancers). Variants within each of these elements may have largely independent effects from one another^49,67^ and from those within protein-coding regions^116^. Furthermore, we identify cases, such as *IKZF1/DDC*, where multiple causal variants in the same LD block intersect the regulatory elements of different candidate genes, leading to their joint prioritisation. These results reinforce the notion that the assumption of a single causal variant per LD block used by many established GWAS analysis methods (particularly those based on summary data) is unnecessarily restrictive and may miss key genetic mechanisms underpinning disease processes.

While the enrichment of GWAS signals within enhancers was first demonstrated over a decade ago^16^, with the first studies leveraging 3D information for enhancer-gene assignment following shortly thereafter^117–119^, the majority of GWAS gene prioritisation studies to date still do not consider 3D chromosomal data^120^. Nonetheless, several computational approaches for variant-to-gene assignment integrating fine-mapped GWAS signals with 3D genomics information and other sources of evidence are now becoming available. For example, FUMA SNP2GENE provides the option to identify candidate genes via enhancer-promoter interactions, but does not integrate fine-mapping SNP probabilities^121^. In addition, the L2G (locus-to-gene) pipeline uses a machine learning algorithm that integrates multiple features, including Capture Hi-C^122^. L2G provides an interpretable output that shows the relative contributions of many factors, including QTL colocalisation, genomic distance, VEP scores^123^, and enhancer-promoter interactions, towards an overall gene score per credible set. L2G is available on the OpenTargets platform^37^, but it is not easily adaptable to new functional data. Finally, H-MAGMA incorporates Hi-C-derived chromatin interactions to refine SNP-to-gene assignment for non-coding GWAS variants, but does not integrate them into a probabilistic framework^124^. MultiCOGS complements these efforts by providing an unsupervised and interpretable Bayesian framework based on cell-type-specific, mechanistically-grounded readouts that can be applied to 3D genomic data in cell types relevant to the disease context.

Using multiCOGS across six autoimmune traits to prioritise disease risk-linked genes with potential roles in ILC3s, we produce a compendium of 251 genes, including both known and potentially novel candidates. Integration with a CRISPRi screen for genes affecting ILC3 inflammatory response provides a first indication of their potential role in ILC3 biology. This includes 11 prioritised genes that were detected as putative IL-22 activators and repressors in the CRISPRi screen^125^. However, further targeted experiments are still required to gain a deeper understanding of the functional role of the prioritised genes in ILC3 biology and their contribution to autoimmune disease risk.

The *Cln3* gene, prioritised in our analysis for CD risk in ILC3s but not in CD4+ T cells, underlies the majority of cases of the neurodevelopmental disorder Batten disease. While immune features have been reported in Batten disease and other lysosomal disorders^126,127^, the function of *Cln3* in the immune system remains poorly understood. Here, we show that *Cln3* expression is downregulated upon cytokine stimulation of mouse ILC3s, and that *Cln3* overexpression in an ILC3-like mouse cell line impacts stimulation-induced transcriptional programmes and cytokine production. In contrast, CRISPRi knockdown of *Cln3* did not show a pronounced phenotype in our model system, and, consistent with this, was not detected as a significant hit in the CRISPRi screen for regulators of ILC3 inflammatory response.^125^ CLN3 is a transmembrane lysosomal protein with established roles in vesicular trafficking and lysosomal homeostasis^128^. Consistent with this biology, our functional data support a role for activation-induced downregulation in promoting the inflammatory capacity of ILC3s. In addition to its trafficking functions^109,110,111^, recent studies have demonstrated that CLN3 is required for the catabolism of glycerophospholipids^87,129^, which are key structural components of cellular membranes and have emerging regulatory roles in innate immune signalling. Accordingly, *Cln3* knockdown in mouse monocytes was shown to interfere with LPS-induced secretion of the inflammatory cytokine IL-6^,130^. These observations raise the possibility that CLN3 may influence immune effector functions through effects on membrane composition, vesicular dynamics, or both. Together, our findings implicate CLN3 in the regulation of ILC3 inflammatory function and CD risk, raising the possibility that inflammatory processes may contribute to gastrointestinal manifestations observed in CLN gene deficiency.^131^

Notably, the region harbouring the fine-mapped CD susceptibility variants in the *CLN3* locus lacks active chromatin signals in ILC3s, as well as in other cell types represented in the Ensembl Regulatory Build. This suggests that regulatory activity at this locus may be highly context-specific, potentially emerging only under inflammatory conditions or within discrete cellular states. Supporting this notion, H3K36me3 deposition across this region in lymphoblastoid cell lines was recently proposed as a mark of enhancers that are ‘poised’ for rapid activation^132^. However, CD-associated variants in this locus may also exert regulatory effects through alternative mechanisms. Several fine-mapped variants in the *CLN3* locus are linked to alternative polyadenylation of the *CLN3* transcript’s 3’UTR across multiple tissues^133,134^, a mechanism that can influence mRNA stability and translational efficiency and is increasingly recognised as a contributor to complex disease risk^134^. In addition, C*LN3* was reported to undergo splicing-dependent transcriptional activation^135^, further expanding the range of potential regulatory mechanisms operating at this locus. The regulatory complexity of the *CLN3* locus is further augmented by its detection as an eQTL for multiple neighbouring genes across diverse cell types. In monocytes, this locus is also an eQTL for the known CD gene *IL27,* with an opposite direction of allelic effect and a lower statistical significance relative to *CLN3* itself^105,136^. Notably, *IL27* is not appreciably expressed in either mouse or human ILC3s. In addition, *CLN3* shares eQTLs with, and is divergently expressed from, the apolipoprotein B receptor gene *APOBR*. Consistent with this, we show that *Cln3* and *Apobr* are co-regulated upon IL-23/IL-1β stimulation in a mouse ILC3-like cell line. APOBR has a recognised role in lipid uptake in myeloid cells^137^, but its function in the lymphoid compartment remains unclear and is likely mechanistically distinct from that of CLN3.

Human ILC3s in our study are derived from tonsillectomy material, but their regulatory elements show an enrichment for variants associated with immunological disorders affecting a broad range of tissues. This is consistent with findings from single cell genomics suggesting that cell type, rather than tissue type, is likely to be the driving factor behind variation in chromatin accessibility and gene expression^138,139^. Furthermore, ILC3s from regularly inflamed tonsils have a closer cytokine profile to mucosal-resident ILC3 populations than ILC3s from resting lymph nodes or peripheral blood^140^. Focused studies in relevant physiological contexts and disease models will further establish the role of ILC3s in mediating the effects of genetic variation. These analyses are, however, complicated by the rarity of ILC3s and a lack of robust human cell line models for this cell type, as well as the strong influence of organismal and environmental factors, which are difficult to reproduce in a laboratory setting either *in vitro* or *in vivo,* on autoimmune disease pathogenesis.

In conclusion, we present updated methodologies for profiling and detecting promoter-anchored interactions and for leveraging these data to interpret GWAS signals. Using this framework, we provide a comprehensive catalogue of regulatory chromatin contacts and candidate autoimmune risk genes in ILC3s, and take initial steps toward their functional validation. These findings advance our understanding of ILC3 biology and the contributions of this rare cell type to disease, and highlight the utility of our approach for dissecting regulatory architecture in other rare cell types and complex traits.

## Methods

### Human ILC3 cell isolation

Three children requiring tonsillectomy were recruited to a prospective study at a tertiary academic care centre through the division of Pediatric Otolaryngology-Head and Neck Surgery at Cincinnati Children’s Hospital Medical Center with an institutional review board (IRB) approval. Criteria for enrollment in the study included a history of sleep-disordered breathing or recurrent or chronic tonsillitis requiring removal of the tonsillar tissue. Consent was obtained from parents in the perioperative suite on the day of the procedure. Subjects were excluded from the study if the tonsillar tissue was acutely infected or if anatomic abnormalities were present requiring a more detailed pathologic evaluation post the surgical procedure. Samples were labelled with a de-identified barcode and transferred to the research team for further processing.

Next, tonsils were dissociated into a single-cell suspension as previously described^141,142^. Briefly, Human tonsil tissue was processed by mincing with scissors, followed by transfer of up to 4g of tissue to a gentleMACS C tube (Miltenyi Biotec) containing 8 mL of phosphate-buffered saline (PBS) with 0.5 mg/mL collagenase D and 3000 U/mL DNase I, then dissociated on a GentleMACS Octo Dissociator (Miltenyi Biotec) using “program C (Spleen program 2 followed by spleen program 3).” Tissue homogenates were incubated in a 37°C water bath for 15 minutes, then dissociated again using “program C” and transferred through a 100 μm cell strainer into 20mL RPMI containing 10% human AB serum (Sigma Aldrich). Next, the cell suspension was overlaid on 10mL of Ficoll-Paque PLUS (GE Healthcare) and subjected to density-gradient separation via centrifugation for 20 min at 1800 rpm, 20°C, slow acceleration and no brake. Leukocytes were collected from the interphase layer and then washed with 50mL of PBS for 6 minutes at 1600 rpm, 20°C.

Single cell suspensions of tonsil mononuclear cells were subjected to positive selection with anti-human-CD3, anti-human-CD19 and anti-human-CD14 (Miltenyi Biotec) and transferred through LD columns (Miltenyi Biotec) according to the manufacturer’s guidelines (**Fig. S9**). The depleted cell suspension flowthrough was collected into a 15mL conical tube and then centrifuged for 5 minutes at 1200rpm, 20°C. Subsequently, cells were labelled with LIVE/DEAD™ Fixable Near-IR dead cell stain kit (Invitrogen). Next, cells were labeled with sorting antibody cocktail which contained negative linage (Lin-) CD19 Brilliant Violet (BV)421 (HIB19), CD14-BV421 (63D3) and CD3-BV421 (OKT3), and the following antibodies: CD45-FITC, (HI30), CD94-PerCP-Cy5.5 (DX22), CD127-PE-Cy7 (A019D5), cKit-BV510 (104D2) and NKp44-Alexa Fluor (AF)647 (P44-8) all purchased from Biolegend (San Diego, CA), CRTH2-PE (301109, R&D). ILC3 cells were sorted based on the expression of CD45+Lin-CD127+CD94-CRTH2-cKit+NKp44+, similarly to Bar-Ephraim et al. Cell sorting was performed using a FACSAria II sorter (BD Biosciences, Mountain View, CA, USA). Post sorting sorted ILC3 cells were washed with PBS for 5 minutes at 1200rpm, 20°C and then incubated in 100 uL of 2% formaldehyde (in PBS) for 10 minutes, followed by the addition of 0.125M glycine. Next, cells were centrifuged at 400g for 5 minutes at 4°C, resuspended with cold PBS and centrifuged again at 400g for 5 minutes at 4°C, supernatant was discarded, and cells were snap-frozen in liquid nitrogen and then stored at −80°C prior to PCHi-C analysis.

### Human CD4+ T cell isolation

Total CD4+ lymphocytes were obtained from PBMCs from venous blood by negative selection using EasySep Human CD4+ T Cell Enrichment kit (Catalog #19052) from STEMCELL Technologies. Purified CD4+ T cells were washed with PBS for 5 minutes at 1200 rpm, 20°C and then incubated in 100 μL of 2% formaldehyde (in PBS) for 10 minutes, followed by the addition of 0.125M glycine. Next, cells were centrifuged at 400g for 5 minutes at 4°C, resuspended with cold PBS and centrifuged again at 400g for 5 minutes at 4°C, supernatant was discarded, and cells were snap-frozen in liquid nitrogen and then stored at −80°C prior to PCHi-C analysis. Two replicates of 1 million and two more replicates of 50,000 cells were used to generate PCHi-C datasets. The samples were obtained from two male donors after written informed consent under studies “A Blueprint of Blood Cells,” REC reference 12/EE/0040, and “Genes and mechanisms in type 1 diabetes in the Cambridge BioResource,” REC reference 05/Q0106/20; both approved by the NRES Committee East of England – Cambridgeshire and Hertfordshire.

### Promoter Capture Hi-C

Promoter Capture Hi-C was performed as previously described^44^. Cells were lysed in a lysis buffer (30 minutes on ice), and digested with *DpnII* (NEB) overnight at 37°C while rotating (950 rpm). Restriction overhangs were filled in with Klenow (NEB) using biotin-14-dATP (Jena Bioscience), and ligation was performed in the ligation buffer for 4 hours at 16°C (T4 DNA ligase; Life Technologies). After overnight de-crosslinking at 65°C, the ligated DNA was tagmented to produce fragments of 300-700 bp. Ligation products were isolated using MyOne C1 streptavidin beads (Life Technologies), followed by washing with Wash&Binding buffer and nuclease-free water. Isolated Hi-C ligation products on the beads were then used directly for PCR amplification, and the final Hi-C library was purified with AMPure XP beads (Beckman Coulter). Promoter Capture Hi-C was performed using a custom-designed Agilent SureSelect system following the manufacturer’s protocol. The PCHi-C libraries were paired-end sequenced (100 bp) on an Illumina HiSeq 2500 machine at a sequencing depth of ∼400 million reads per sample (**Table S1**).

### PCHi-C data pre-processing and detection of significant interactions

Sequencing data from three ILC3 PCHi-C biological replicates were aligned to the hg38 genome assembly using Bowtie2^143^ and quality-controlled using HiCUP^144^. Quality metrics for all generated PCHi-C datasets are reported in **Table S1**. Significant interactions were then detected across the replicates by CHiCAGO^39^ as previously described^40^ at single *DpnII* fragment resolution and in bins of fragments approximately 5 kb in length, with the baited promoter fragments left solitary (unbinned).

Leaving the baited *DpnII* fragment unbinned meant that nearly every baited fragment was occupied by a single protein-coding gene promoter. In contrast, a third (33%) of baited fragments in the *HindIII*-based Capture Hi-C design (with a median fragment size of 4 kb) contained two or more promoters. Therefore, leaving the baited fragment unbinned significantly improved the resolution and interpretability of analyses such as (multi)COGS.

For CHiCAGO analysis at single-fragment resolution, p-value weights were estimated following our previously described procedure^40^ and are listed in **Table S11**; default p-value weights were used for the 5 kb analysis. A CHiCAGO score cutoff of ≥5 was used for both resolutions. A consensus list of promoter interactions was compiled from non-redundant contacts detected at the fragment and 5 kb resolutions.

### Integration with *HindIII* Promoter Capture Hi-C data

Our previous PCHi-C study in 17 abundant human primary blood cell types, including both lymphoid and myeloid cells^30^ was performed using a 6 bp restriction enzyme *HindIII*, unlike the 4-bp cutter enzyme *DpnII* used in the current study. Since restriction fragment size affects the distance distribution of contacts detected in Hi-C-related methods^40,145,146^, direct comparison across these two datasets is challenging. To partially address this issue, we pooled the reads in the *DpnII*-based ILC3 data into genomic windows corresponding to *HindIII* fragments and re-processed the data with HiCUP using the hg19 genome assembly and *HindIII* parameters. We then identified significant interactions using CHiCAGO^39^ with the default *HindIII-*based parameters and integrated them with the significant interactions from the Javierre et al. study^30^. To assess the similarity of promoter-interaction patterns in ILC3s with the cell types profiled in Javierre et al., we first ran a joint PCA analysis. We noted that PC1 (accounting for <10% of the variance) clearly segregated the three ILC3 replicates from the remaining cell types, and therefore most likely corresponded to the difference in PCHi-C methods, resolution and sequencing depth. We disregarded PC1 and focused on PC2, PC3, and PC4, accounting for 6.16%, 3.7%, and 3.16% of variance across all tissues, respectively (components beyond PC4 accounted for <3.1% of variance each and were disregarded). For visualisation purposes, we combined these three components using the UMAP non-linear dimensionality reduction algorithm implemented in the umap package in R^147^, obtaining the plot shown in **Fig. S1A.**

### Alternative promoter analysis

We used the CHiCAGO results for ILC3 PCHi-C data at 5 kb resolution to profile PIR sharing between alternative promoters. First, we identified a set of genes that had more than one baited promoter, with each promoter having at least one significant interaction with a CHiCAGO score of ≥5 with ≥5 reads. We defined fully shared PIRs as those that interacted with all baited alternative promoters for the same gene, and partially shared PIRs as those that interacted with a subset of alternative promoters for the same gene. We defined distinct PIRs as those that only interacted with a single promoter fragment (CHiCAGO score ≥5). To increase the stringency with which we called PIRs “distinct”, we applied two further criteria. First, if a PIR interacted with another alternative promoter at a lenient CHiCAGO score ≥3, we defined that PIR as shared. Second, if the adjacent fragment to the PIR in question interacted with another alternative promoter at a CHiCAGO score ≥3, we also defined that PIR as shared. We note that, under our classification rules, the PIRs of genes with only two alternative promoters included in the analysis can only be classified as “fully shared” or “distinct”. Therefore, the “partially shared” PIR category was only applicable to the subset of genes with more than two baited alternative promoters.

### Epigenomic data pre-processing

For epigenetic data analysis in ILC3s, the SRA accession list was downloaded from the GEO accession GSE77299. The SRA files were converted to FASTQ file,s and sequencing adapters were trimmed from reads using *trim galore* (https://github.com/FelixKrueger/TrimGalore). The reads were filtered by PHRED score ≥30 and examined for proper pairing with a mate (when paired-end). The sequencing quality and duplication level were checked using FastQC (https://www.bioinformatics.babraham.ac.uk/projects/fastqc/). Sequences were mapped to the hg38 reference genome using STAR with modifications for aligning ChIP-seq and ATAC-seq reads. Samtools^148^ was used to select reads with a MAPQ score of 255, which is the flag for uniquely mapping reads from STAR^149^. ATAC-seq reads were filtered, retaining properly paired and oriented reads using the samflag=3. PCR duplicates were removed using samtools. We then removed reads that fell within blacklisted regions using Bedtools^150^ intersect. The final filtered BAM file was then converted to a BED file using Bedtools bamtobed. This conversion breaks read-pairing and ensures each read contributes to peak identification with MACS2^151^. The ATAC-seq reads in BED format were shifted by +4 bp on the (+) strand, and −5 bp on the (-) strand to account for the Tn5 transposase cut site. Peaks were called using MACS2 using three biological replicates per sample as the treatment group with an input ChIP-seq control sample. The replicate correlation between the ATAC-seq samples was poor, with a <10% overlap between biological replicates. This result was consistent with the high level of duplication and low peak count (8,852) in the worst sample (SRR3129112). Thus, our ATAC-seq results were limited to the sample withthe best quality metrics (SRR3129113). In total, we detected 34,077 H3K27ac peaks and 72,825 ATAC-seq peaks. For epigenetic data analysis in CD4+ T cells, we used BLUEPRINT epigenome datasets from male donors C002Q1, S008H1, and S007G7.

### Activity-By-Captured-Contact (ABCC)

For a given gene-enhancer pair, the ABC score is the normalised product of enhancer Activity (proxied by the levels of chromatin accessibility and relevant histone modifications) and Contact (proxied by 3D contact frequency detected from a chromosome conformation capture assay)^14,41^. In the original implementation of ABC, Activity is estimated as the geometric mean of read counts of DHS/ATAC-seq peaks and Contact by KR-normalised Hi-C contact frequency between the respective element and gene promoter^41^. The resulting product is divided by the sum of all ABC values for a given gene from enhancers within a 5-megabase window around the transcription start site:

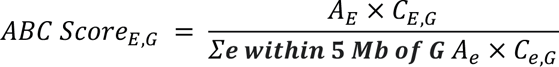

To adapt ABC for PCHi-C data, we took advantage of the CHiCAGO normalisation algorithm and developed an imputation procedure in the normalised counts space based on the inferred decay of interaction read counts with distance. As we do not expect the frequency of enhancer-promoter contacts to fall below levels expected due to Brownian collision, for a given pair of fragments involving a baited promoter, we selected the maximum between the CHiCAGO-normalised observed read counts (N_obs_) and expected read counts N_exp_ estimated as:

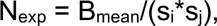

where B_mean_ is the CHiCAGO-estimated Brownian noise level and s_i_ and s,_j_ and the bait- and other end-specific scaling factors. For promoters that could not be baited in the Capture Hi-C design and those that were filtered out due to QC fail, we estimated the expected normalised read count directly from the interaction distance *d*, using the distance function f(*d*) fitted by CHiCAGO. Due to the strong bias of the distance function *d* towards the very short range interactions (<1.5kbp) and to ensure we do not disregard long-distance interactions, in the imputation procedure we introduced a contact frequency cap for candidate enhancers that are closer than at least one fragment away from the bait equal to the contact frequency prediction of distance function *d* at 1.5 kbp (median fragment length). Please refer to Additional File 1 in the publication presenting the CHiCAGO pipeline^39^ for the formal definition of these parameters and their estimation procedures.

The imputed normalised read counts were used as Contact data in the ABC pipeline, and the public H3K27ac and ATAC-seq data in ILC3s processed as described above were used to compute Activity. To validate the ABCC approach, we took advantage of the high-throughput CRISPRi-FlowFISH data from Fulco et al.^41^, which presented the impact of perturbing ∼3,500 enhancer elements on the expression of 30 genes in K562 cells. Since PCHi-C data for K562 cells are not currently available, we used our previously published PCHi-C dataset in the related primary cell type, erythroblasts^30^, to generate the ABC scores based on these data and the ATAC-Seq and H3K27ac ChIP-Seq datasets for K562 cells from Fulco et al. In comparison with the original ABC scores from Fulco et al. based on pooling conventional Hi-C data from multiple cell types, our approach showed a higher precision (69.1% vs 58.3%) at the same level of recall (58.3%) of CRISPRi-FlowFISH-validated enhancer-promoter pairs (**Fig. S2**). To select ABCC score cutoff, we optimised the Pearson correlation between per-gene ABCC numerator and gene expression (*R_ABC-GE_*), in an approach inspired by Xu et al.^152^. We opted to use a single ABCC score cutoff of 0.023 in all analysed cell types, as it was close to the maximum *R_ABC-GE_* in each cell type, as well as to the cutoff of 0.02 that yielded an optimal precision-recall of CRISPRi-FlowFISH-validated enhancer-promoter pairs in K562 cells.

### Microarray gene expression data analysis

The microarray CEL files were downloaded from the GEO accession number GSE78896. The CEL files were then analysed using AltAnalyze (http://www.altanalyze.org/). Probes were filtered for a DABG (detection above background) as previously described^153^. Probes were collapsed to the gene level and RMA-normalised using the AltAnalyze platform.

### RNA-seq data analysis

Human ILC3 RNA-seq data were downloaded from the GEO accession number GSE130775. Salmon^154^ was used to quasi-map reads to transcripts. Reads were aligned to the hg38 genome assembly. The transcript counts were then imported and collapsed to gene counts using Tx import.

Mouse ILC3 differential RNA-seq data analysis was performed using DESeq2^155^. In brief, the gene count matrices were downloaded from GEO (GSE120723) and the standard DESeq2 algorithm was run according to the vignette. Low-count genes were pre-filtered before running. The following parameters were used to report significantly differentially expressed genes: alpha = 0.05 and adjusted p-value < 0.05.

### PIR enrichment for epigenomic features

For each gene, sets of adjacent PIRs for each gene (detected at the fragment or 5 kb resolution or the merged PIR sets for each gene) were collapsed together to obtain “collapsed PIRs” (cPIRs). Trans-chromosomal PIRs were removed. The observed proportion of cPIRs overlapping epigenomic features of interest (ATAC-seq, H3K27ac or H3K4me3, respectively) was computed using the *foverlaps* function from the *data.table* package in R. To obtain the expectation for this proportion, we repeated this analysis for random cPIRs that were generated by “transplanting” each set of all cPIRs for each gene to randomly selected genes in a manner preserving the size and spatial localisation of the cPIRs with respect to each other and the respective baited promoter fragment. This “transplantation” was repeated 100 times for all genes (baited promoter fragments), and the mean proportion of random cPIRs overlapping epigenomic features of interest (over 100 permutations), as well as the standard deviation of this quantity, were compared with the proportion of overlap for the observed cPIRs. Compared with the PIR enrichment estimation algorithm implemented in CHiCAGO (*peakEnrichment4Features*), this permutation procedure preserves not only each PIR’s distance from bait, but also the spatial relationships between multiple PIRs of the same gene.

### LOLA enrichment analysis

We performed LOLA v1.18^156^ enrichment analysis to assess whether active and/or open regulatory elements of multiCOGS-prioritised genes were enriched for specific transcription factor binding sites and chromatin features compared to all genes tested by multiCOGS.

We defined active/open PIRs as those with overlapping ATAC-seq or H3K27ac ChIP-seq peaks within significant PIRs identified by promoter capture Hi-C interactions (CHiCAGO) or predicted by our ABCC algorithm for multiCOGS-prioritised genes. The background universe comprised all active/open PIRs from the same datasets for all tested genes. Regions were converted to GRanges objects using the GenomicRanges package, and enrichment was tested using the LOLA core pipeline with the LOLA Core RegionDB, using default parameters. Significant enrichments were defined as those with q-value < 0.05.

### RELI analysis

RELI^56^ (v0.1.1a) was used to find enrichment of genetic variants in promoter-interacting regions (PIRs) that are accessible and marked with activating epigenetic markers (H3K27ac and ATAC-seq). In brief, RELI tests genomic features such as ATAC-seq, ChIP-seq, or PIRs for statistically significant overlaps with known disease risk variants identified from genome-wide association studies. Risk variants are expanded to linkage disequilibrium blocks (LD blocks) with variants that have an R^2^ value ≥ 0.8. LD blocks are then intersected with the genetic feature BED files. A null distribution is generated using randomly shuffled LD blocks (n=1,000) and performing the intersection with the feature files. A p-value is generated by comparing the observed number of intersections in the test to the null distribution.

Promoter-interacting regulatory elements were determined as input for RELI as follows. The PIR sets were the union of PCHi-C interactions (CHiCAGO score ≥ 5, binned to 5 kb or DpnII fragment-level resolution) and ABC enhancers, excluding any trans-chromosomal interactions. Regulatory elements were then defined as the union of peaks of open chromatin and H3K27ac in ILC3 and CD4+ T cells (using ATAC-seq and ChIP-seq data as above). The true intersection between these regulatory elements and PIRs in each cell type was then determined using pybedtools *intersect*. The coordinates for these regions were lifted over from hg38 to hg19 using UCSC liftOver (v. 377), then sorted and merged for use with RELI. RELI was run against all 495 traits with ≥ 10 independent risk loci and of European ancestry in the GWAS Catalog. Bonferroni and Benjamini-Hochberg p-value correction were performed with the Python package statsmodels, with alpha=0.05 (family-wise error rate of 5%; the probability that at least one of the predictions is a false positive). Traits with the BH-adjusted p-value < 0.05 were defined as significant. For depicting RELI results, we labelled only significant traits with enrichment ≥2.

### Standard COGS

To run standard COGS^30,31^, we adapted the code from the R package rCOGS (https://github.com/ollyburren/rCOGS) to use the *data.table* framework instead of *GenomicRanges* for optimised speed and to enable both the standard COGS and multiCOGS analyses (see Code availability). We used linkage disequilibrium blocks calculated for GRCh38 from https://github.com/jmacdon/LDblocks_GRCh38^157^ and minor allele frequencies from the 1000 Genomes Project, European individuals. Protein-coding SNPs were identified using VEP version 99.2 (https://github.com/Ensembl/ensembl-vep). We obtained gene transcription start sites (Havana and Ensembl/Havana merge) from Ensembl GRCh38 release 88 (March 2017), matching the version used to design the *DpnII* promoter capture system. We included promoters irrespective of whether they were targeted in the capture system, enabling COGS to prioritise all gene targets where the causal variants fell near the gene promoter (defined as +/− 5 *DpnII* fragments from the transcription start site). PIRs with CHiCAGO interaction scores ≥5 or ABC scores of ≥0.04 were used as COGS input. The results for each protein-coding gene were linked across datasets using Ensembl gene IDs as primary identifiers. The Major Histocompatibility Complex was removed (GRCh38 6:28510120-33480577) prior to running COGS.

### Sources of prior mechanistic evidence for CD genes

Datasets used to compare the COGS prioritised genes with previously functionally validated genes were: OpenTargets^37^ (L2G gene prioritisation score > 0.5 for five CD studies^71,158–161)^, the IBDDB database of functionally validated targets^80^, a functional screen of IBD genes^81^, experimentally validated IBD and CD genes from DisGeNET^82^ that had evidence “AlteredExpression”, “Biomarker”, “Posttranslationalmodification”, or “Therapeutic” or CD-containing exonic variants in a recent IBD exome study^83^.

### Multivariate GWAS fine-mapping

The Sum of Single Effects (SuSIE) model allows for multiple causal variants within a GWAS locus^68,69^. We downloaded summary data for Crohn’s disease^71^ (GCST004132), Ulcerative colitis^71^ (GCST004133), Inflammatory Bowel Disease^71^ (GCST004131), Celiac Disease^162^ (GCST000612), Adult onset Asthma^163^ (GCST007799) and Primary Sclerosing Cholangitis^164^ (GCST004030) from the GWAS Catalog. and used LD block data for EUR from lddetect (https://bitbucket.org/nygcresearch/ldetect-data/src/master/), which we liftOvered^165^ to hg38 to divide the data into approximately independent blocks. We used EUR samples from phased 1000 Genomes Phase 3 data, downloaded from https://mathgen.stats.ox.ac.uk/impute/1000GP_Phase3.html, to generate LD matrices. We used these matrices to first impute the summary statistic data within blocks using the published method^66^. For blocks with appreciable association signals (minimum p < 10^-6^), we used the susieR package^68,69^ to fine-map the data. We defined “detected signals” as those for which SuSiE could calculate a 95% credible set, and used the posterior inclusion probabilities (PIP) for each SNP for each signal thus detected as input for multiCOGS, described below. For the remaining blocks, or where susieR failed to find any signals meeting our criteria, we fine-mapped using the single causal variable approach, as previously described^30,31^, and used the posterior probabilities of association as input for multiCOGS.

### multiCOGS

We modified the COGS algorithm to account for the inclusion of multiple association signals in a region (“multiCOGS”). While in standard COGS, fragment-level scores are calculated by summing variant-level posterior inclusion probabilities (PIP, calculated as above) within a given fragment and LD block, multiCOGS considers each credible set within each LD block and forms an overall gene score as probability that at least one of the multiple fine-mapped signals is linked, through PCHi-C, to the gene of interest:

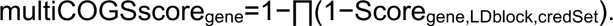

To reveal the contributions of the four categories of genomic loci underlying the prioritised genes (PCHi-C PIRs, ABC enhancers, promoter proximal regions and coding SNPs) we additionally ran multiCOGS on each category separately by specifying the *feature.names* argument in the *compute_cogs* function.

### Assessing the biological function of CD-prioritised genes

The Gene2Func tool in FUMA (v1.5.2) was run using all multiCOGS genes with a score ≥0.5, Ensembl version 102, and GTex v8. As a background, we used all genes with assigned multiCOGS scores in ILC3s, of which 17,984 had a recognised Ensembl Gene ID in FUMA. Multiple testing correction was done via the Benjamini-Hochberg method (FDR) with an adjusted p-value cutoff of 0.05 and a minimum of 2 genes in a set. The MsigDB version was v7.0. We additionally checked for enrichment of multiCOGS genes in The Inflammatory Bowel Disease Transcriptome and Metatranscriptome Meta-Analysis (IBD TaMMA) Framework^91^. We filtered the 496 datasets of differentially expressed (DE) genes (adjusted p-value < 0.05 and absolute log2 fold change ≥2) that were compared across the same tissues and selected only sets with a maximum of 2,000 DE genes, to avoid mis-estimation of the normalised enrichment score, resulting in 24 datasets. Then we ran the *enricher* function in the R package clusterProfiler^166^ (version 4.2.2) for all multiCOGS genes with a score ≥0.5.

### Cell culture

Mouse MNK-3 cells^112^ and the derived lines were cultured in DMEM with glucose/pyruvate/ L-glutamine supplemented with 10% fetal bovine serum, 1X penicillin-streptomycin, 10 ng/ml mouse recombinant IL-2 and IL-7 (R&D Systems), and 50 µM 2-mercaptoethanol. Media for CRISPRi MNK-3 (MNK-3i) cells contained 10 µg/ml blasticidin S, and media for CRISPRa MNK-3 (MNK-3a) cells contained 10 µg/ml blasticidin S and 1250 µg/ml hygromycin B. MNK-3i/a cells with sgRNA additionally received 2 µg/ml puromycin. MNK-3 activation was induced with 10 ng/ml IL-1β and 10 ng/ml IL-23 (R&D Systems).

### CRISPR activation and interference

MNK-3i cells were generated as described^167^ from parental MNK-3 cells. In brief, MNK-3 cells were transduced with lentivirus containing pLenti CMV rtTA3 Blast (Addgene #26429), selected by blasticidin S, and then infected with TRE3G-dCas9-KRAB-P2A-mCherry lentivirus. Following incubation with doxycycline, mCherry-positive cells were subcloned, and Western blot analysis confirmed robust expression of doxycycline-inducible dCas9-KRAB. MNK-3a cells were lentivirally engineered from MNK-3 to constitutively express the dCas9-VP64 fusion gene (Addgene #61425) and the MS2-p65-HSF1 transactivator complex (Addgene #89308), selected by blasticidin S and hygromycin B, and subcloned. All cells were tested for mycoplasma.

Sequences for Cln3-targeting and scrambled gRNAs were based on published sgRNA libraries for MNK-3i^168^ and MNK-3a^169^ are listed in **Table S14** alongside RT-qPCR primer sequences. sgRNA sequences and their reverse complement were synthesised by Sigma, annealed, and cloned into lenti sgRNA(MS2)_puro optimised backbone (Addgene #73797) for MNK-3a or sgOpti (Addgene #85681) for MNK-3i using Esp3I digestion as previously described^170^. sgRNA plasmid integration was confirmed by Sanger Sequencing (Ohio State Comprehensive Cancer Center Genomics Core, Columbus, OH, USA). Lentiviral plasmids pMD2.G (Addgene #12259) and psPAX2 (Addgene #12260) were transfected along with the sgRNA plasmid into HEK293T cells (Mirus TransIT-293T transfection reagent). Lentivirus media was harvested and filtered 48-72 hr post-transfection. Puromycin selection began 36 hr after lentiviral guide transduction into MNK-3i/a cells in the presence of polybrene. Bulk transduced populations were used for experiments and maintained in selection antibiotics. RT-qPCR confirmed repression (MNK-3i lines after 48 hr doxycycline incubation) or overexpression (MNK-3a) of target genes relative to *Actb* and respective scramble control (Trizol RNA isolation; Verso cDNA synthesis).

To induce CRISPRi guide expression, MNK-3i stably expressing Cln3-targeting and scrambled (Scr) gRNAs were incubated with 2 µg/ml doxycycline for 48 hr. To confirm stimulation, cells were harvested 21 hr after cytokine stimulation and stained for intracellular IL-17F and IL-22 (eBioscience IL-22 clone 1H8PWSR and IL-17F clone eBio18F10; BD Life Sciences Cytofix/Cytoperm kit). Expression of IL-17F and IL-22 was assessed on FACSymphony (BD Life Sciences) and compared against a respective scrambled control.

### RNA-sequencing

RNA was harvested by spin column (Qiagen RNeasy kit) for polyA-selected 2×150bp bulk RNAseq (Illumina platform, University of Cincinnati Genomics, Epigenomics, and Sequencing Core, Cincinnati, OH, USA). RNA-seq samples were generated in triplicate.

Raw paired-end RNA-seq reads were quantified using *kallisto* (v0.48.0) against the mouse reference transcriptome (GENCODE release M32, GRCm39). Transcript indices were first generated with *kallisto index*, and transcript abundances were quantified for each sample using *kallisto quant* with 100 bootstrap replicates. Transcript-level abundance estimates were subsequently summarised to the gene level in R using the *tximport* package (v1.30.0) together with a transcript-to-gene mapping file. Sample metadata, including experimental condition, CRISPR status, and replicate information, were compiled into a metadata table. Gene-level count matrices generated by *tximport* were then used as input for normalisation and differential expression analysis with *DESeq2* (v1.38.0). Sample metadata, including experimental condition, CRISPR status, stimulation, and replicate information, were compiled into a metadata table.

Gene-level count matrices were then used for normalisation and differential expression analysis with DESeq2 (v1.38.0). A variance-stabilising transformation (rlog) was applied for visualisation and principal component analysis to identify batch effects. Differential expression analyses were performed using linear models incorporating relevant covariates. For wild-type samples, stimulation status was tested while including CRISPR type as a batch covariate. For CRISPRa and CRISPRi samples, models including interaction terms between CRISPR treatment and stimulation were used to assess treatment-specific effects. Adjusted p-values were calculated using the Benjamini-Hochberg method, and genes with adjusted p-values < 0.05 were considered statistically significant.

### RNA isolation and quantitative RT–PCR

Total RNA was isolated from snap-frozen cells using QIAshredder columns and the RNeasy spin-column system (QIAGEN). Complementary DNA (cDNA) was synthesised using the High-Capacity cDNA Reverse Transcription Kit (Thermo Fisher Scientific).

Quantitative PCR was performed using TaqMan chemistry with TaqMan Fast Advanced Master Mix (Thermo Fisher Scientific) on a QuantStudio 5 Real-Time PCR System (Thermo Fisher Scientific). Cln3 expression was quantified using the TaqMan Gene Expression Assay Mm00487021_m1 and normalised to the housekeeping gene Hprt using assay Mm03024075_m1. Reactions were performed in technical triplicate. Relative gene expression was calculated using the ΔΔCt method, with MNK-3 cells electroporated with GFP mRNA used as the reference control condition.

### Design and generation of in vitro-transcribed mRNA

The protein-coding sequence of mouse *Cln3* was based on the longest annotated transcript (NM_001146311.3 / ENSMUST00000084589.11). A Myc epitope tag was inserted near the N terminus, between amino acid residues 3 and 4, within a predicted disordered and cytoplasmic region of the protein. The resulting coding sequence was synthesized and used for in vitro transcription by ApexBio.

In vitro–transcribed mRNA was generated with a Cap 1 structure and incorporated N1-methylpseudouridine. Transcripts contained a poly(A) tail and were supplied in RNase-free sodium citrate buffer (pH 6.4) at a concentration of 1 mg ml⁻¹. Control mRNA encoding GFP was generated using the same chemistry.

### mRNA electroporation and cytokine stimulation

MNK-3 cells were electroporated with IVT mRNA using the ATx electroporation system (MaxCyte). 1.0×10⁷ cells were electroporated in a 100 µl reaction containing 20 µg of GFP or myc-tagged Cln3 mRNA (2 µg per 10⁶ cells) using the “Optimization 8” program. Following electroporation, cells were rested for 15 min at 37 °C and then incubated for 15 min at 37 °C in pre-warmed medium supplemented with 10 µg/mL DNase I (Thermo Fisher Scientific), 5mM MgCl₂, and 1 mM CaCl₂ before transfer to complete MNK-3 culture medium.

At 24 hr post-electroporation, cells were seeded at 3.0×10⁵ cells per well in 24-well plates. Transfected cells were cultured for an additional 24 hr in the presence or absence of recombinant mouse 10 ng/mL IL-1β and 10 ng/mL IL-23 (R&D Systems). At 48 hr post-electroporation, supernatants were collected, clarified by centrifugation, and stored at −20 °C. Viable cell numbers were determined by trypan blue exclusion.

### ELISA assay

Cytokines in cell culture supernatants were quantified by ELISA using DuoSet kits for mouse IL-17, IL-22, and GM-CSF (R&D Systems) according to the manufacturer’s instructions. When necessary, samples were diluted to fall within the dynamic range of the standard curve. Absorbance was measured at 450 nm with wavelength correction at 560 nm using a GloMax Discover microplate reader (Promega). Cytokine concentrations were determined by interpolation from standard curves using a four-parameter logistic fit.

Data were analysed using GraphPad Prism. Statistical significance was assessed using unpaired Welch’s t-tests (single experiment) or linear mixed-effects models with genotype as a fixed effect and experiment as a random effect (multiple experiments).

### Immunoprecipitation and immunoblotting

MNK-3 cells were electroporated with GFP or myc-tagged Cln3 mRNA as described above and harvested 24 hr later. Cells were lysed in a non-denaturing buffer containing 50 mM Tris-HCl, 150 mM NaCl, 1 mM EDTA, 1% n-dodecyl-β-D-maltoside (DDM), 10% glycerol, and protease phosphatase inhibitors (Thermo Fisher Scientific). Lysates were clarified by centrifugation at 4 °C.

Myc-tagged proteins were enriched by incubation of clarified lysates with Myc-Trap agarose beads (ChromoTek) for 1 hr at 4 °C with rotation. Beads were washed in buffer containing 0.05% DDM, and bound proteins were recovered for analysis. Input, unbound, and bound fractions were quantified by BCA assay (Thermo Fisher Scientific), denatured in LDS sample buffer with reducing agent, and resolved by SDS–PAGE on 4–12% Bis-Tris gels (Thermo Fisher Scientific). Proteins were transferred to PVDF membranes, stained with Revert 700 Total Protein Stain (LI-COR), and imaged prior to immunoblotting.

Membranes were blocked and probed with antibodies against myc tag (Cell Signaling Technology #2278, 1:1000) or GFP (Invitrogen #A-11122, 1:2000). Fluorescent secondary antibodies were used at 1:10,000 and blots were imaged using the Odyssey DLx Imaging System (LI-COR).

### Querying a CRISPRi screen for regulators of ILC3 inflammatory response for multiCOGS-prioritised genes

The analysis is based on data from Table S5 in Brown et al^125^, containing a gene-level analysis of a CRISPRi screen in MNK-3i cells. In the experiments performed by Brown et al., MNK-3i cells were induced with doxycycline to express CRISPRi (dCas9-KRAB) machinery and were transduced with a lentiviral gRNA library targeting 20,003 genes. The cells were then stimulated by IL-23 and IL-1β and sorted into subpopulations expressing high and negative levels of the inflammatory cytokine IL-22 released by activated ILC3s. The quantity of each sgRNA in IL22^Neg^ and IL22^High^ cells was detected through PCR amplification and next-generation sequencing. To focus on sgRNA targeting expressed genes, the genes were filtered to those with an average transcript per million (TPM) of ≥2.5 in RNAseq data from MNK-3i+scramble (sgSCR) cells treated with dox (48 hr) and stimulated with 10 ng/ml IL-1β/23 (21 hr). The “test” command from MAGeCK (version 0.5.9.5) ^171^ was applied to generate normalised (method = total) gene-level rankings using Robust Rank Aggregation (RRA). The sgRNA enriched in the IL22^Neg^ population pointed towards genes positively regulating IL-22 production, implicating them in ILC3 inflammatory response. In contrast, sgRNA showing enrichment in the IL-22^High^ population points to ILC3 ‘anti-inflammatory’ genes.

In the present study, we first filtered the genes in **Table S5** from in Brown et al^125^ to those that had been profiled in the multiCOGS experiment, based on an identical gene name between the mouse and human data, leading to a total set of 6438 genes. The genes were ranked based on their MAGeCK score for positive or negative regulation of IL-22 production. We then ran GSEA against each of these rankings, for the 142 multiCOGS genes for inflammatory traits, using the “pathway” function in MAGeCK. We considered significant CRISPRi genes to be those with an adjusted p-value < 0.05 in the gene-level RRA analysis.

## Data availability

Raw PCHi-C data generated in this study for ILC3s are deposited in the Gene Expression Omnibus (GEO) under the accession number GSE216267. Processed R data files containing CHiCAGO scores at the fragment-level and 5kb-binned resolution can be found in the same repository. PCHi-C data for CD4+ T cells were deposited to the European Genome-Phenome Archive (EGA) under managed access in accordance with the conditions of donor consent, under the accession number EGAS50000001316. Raw RNA-seq reads and counts for the CLN3 CRISPRi/a experiments in MNK-3 cells are deposited in GEO under the accession number GSE313942. Supplementary Data files, including significant CHiCAGO interactions at fragment-level and 5kb resolution in ILC3 and CD4+ T cells, ABCC pairs in both cell types and DESeq2 objects for the CLN3 CRISPRi/a experiments, were deposited to Open Science Framework (https://osf.io/aq9fb).

## Code availability

Most scripts for analyses used in the paper are available at https://github.com/vmalysheva/ILC3 and https://github.com/malyshevalab/hILCs_CHi-C, with the following exceptions: CHiC-ABC (https://github.com/pavarte/PCHIC-ABC-Prediction), RELI (https://github.com/tacazares/spivakov_pchic_ILC_CD4), SuSiE (https://github.com/chr1swallace/cd-finemapping-scripts), COGS and multiCOGS (https://github.com/FunctionalGeneControl/multiCOGS).

## Contributions

Conceptualisation: V.M., H.R.-J., C.W., S.W., and M.S. Data curation: V.M., H.R.-J., P.A., J.A.W., X.C., S.P., and M.S. Formal analysis: V.M., H.R.-J., N.L., T.A.C., P.A., J.A.W., Z.F.Y., X.C., S.P., C.W., and M.S. Funding acquisition: V.M., P.F., E.R.M., S.W., and M.S. Investigation: V.M., H.R.-J., N.L., R.B., T.A.C., O.C., D.O., P.A., J.A.W., C.P., J.B., X.C., S.P., N.P., C.W., S.W., and M.S. Methodology: V.M., H.R.-J., N.L., R.B., M.D.R., C.P., J.I.J.D., W.R.O., T.N., P.F., S.S., M.T.W., L.C.K., C.W., S.W., and M.S. Project administration: V.M., H.R.-J., and M.S. Resources: R.B., T.A.C., C.B., X.C., S.P., A.W.D., A.S., F.B., M.F., D.F.S., N.P., J.M.W., E.M.O., C.W., and M.S. Software: H.R.-J., P.A., J.I.J.D., M.T.W., L.C.K., C.W., and M.S. Supervision: V.M., H.R.-J., P.A., M.F., M.T.W., L.C.K., J.M.W., E.M.O., C.W., E.R.M., S.W., and M.S. Validation: H.R.-J., N.L., and R.B. Visualization: V.M., H.R.-J., N.L., P.A., and M.S. Writing – original draft: V.M., H.R.-J., and M.S. Writing – review & editing: V.M., H.R.-J., N.L., S.W., and M.S., with contributions from all authors. Joint first authors: V.M., H.R.-J. and N.L. Joint second authors: R.B., T.A.C., O.C., D.O, and P.A. Joint principal supervisors: C.W., E.R.M., S.W., and M.S.

## Acknowledgements

The authors would like to thank Laurence Game and Ivan Andrew at the LMS Genomics facility for sequencing, Michiel Thiecke and Oliver S. Burren for technical advice and Wing Leung for technical assistance. M.S. is core-funded by the Medical Research Council as an MRC Programme Leader (MC-A652-5QA20). C.W. is funded by the Wellcome Trust (WT220788), the MRC (MC_UU_00002/4), GSK and MSD, and supported by the NIHR Cambridge BRC (BRC-1215-20014). Research activities related to this manuscript that were conducted at Cincinnati Children’s Hospital Medical Center were supported by the NIH (U01AI150748 to E.R.M, M.T.W, L.C.K; R01AI153442 to E.R.M; R21AI156185 to E.R.M; R01HG010730, U01AI130830, R01NS099068, R01GM055479, P01AI150585, R01AI141569 to M.T.W; R01AI024717 to M.T.W and L.C.K; R01AR073228 to M.T.W, L.C.K and S.W; R01DK107502, R01AI148276, U19AI070235, U01HG011172, and P30AR070549 to L.C.K); Cincinnati Children’s Research Foundation (ARC Award to E.R.M, M.T.W and L.C.K; Center for Pediatric Genomics grants to E.R.M, M.T.W and L.C.K); and the L.B. Research and Education Foundation (to N.L.). M.T.W. S.W. and E.R.M. are partially supported by the Pilot Grant from the Cincinnati Digestive Health Center (P30 DK078392). O.C. was supported by an NIH training grant T32 AR069512. D.F.S acknowledges the support by NIH grant 5K08HL148551-02. D.O. was supported by an American Heart Association fellowship. Research activities related to this manuscript that were conducted at Ohio State University’s Medical Center were supported by the NIH (R01AI134035 to E.M.O), Pelotonia Scholars Graduate (to R.A.B.) and PostDoctoral (to A.S.) Fellowships. This project was funded in part by a Commercialisation Award from the Babraham Institute to M.S. and P.F and by the VIB core funding to V.M. M.F. was supported by the British Heart Foundation (BHF; FS/18/53/33863), the BHF Cambridge Centre for Research Excellence (RE/18/1/34212) and the NIHR Exeter Biomedical Research Centre. We thank the UK’s National Institute for Health and Care Research (NIHR) BioResource volunteers for their participation and gratefully acknowledge NIHR BioResource centres. The views expressed are those of the author(s) and not necessarily those of the NHS, the NIHR or the Department of Health and Social Care. For the purpose of Open Access, the authors have applied a CC-BY public copyright licence to any Author Accepted Manuscript version arising from this submission.

## Supplementary Figures

**Figure S1.**
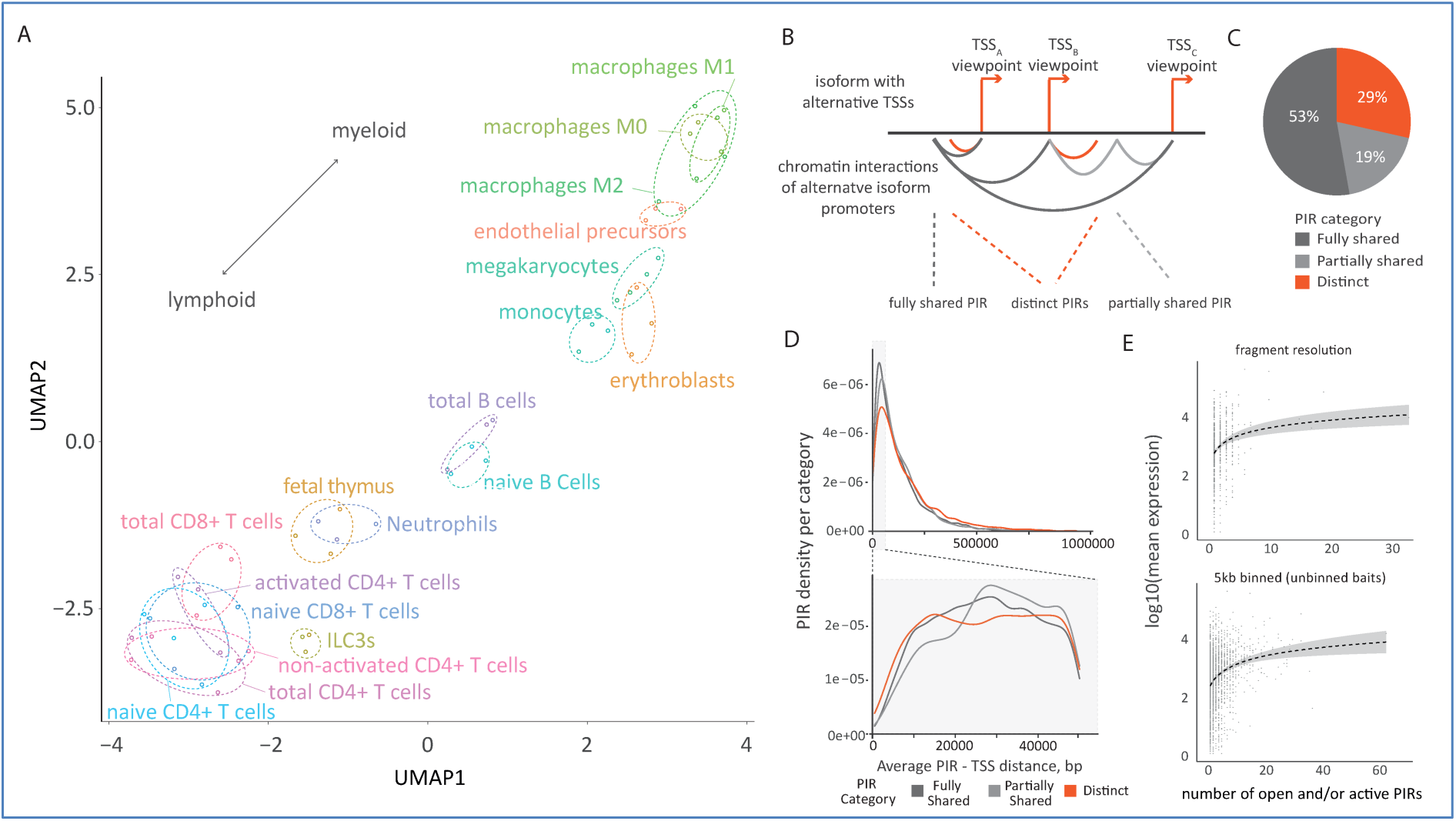
Compendium of promoter-enhancer interactions in ILC3s. **A.** UMAP of CHiCAGO scores detected for PCHi-C in ILC3s versus public data in 17 primary human blood cell types^30^. **B.** Scheme representing the classification of PIRs detected at alternative transcription start sites (ATSS) of the same gene: ‘fully shared’ (shared across all captured ATSSs), partially shared and distinct (unique to a single ATSS). **C.** Pie chart showing the degree of enhancer sharing across alternative transcription start sites (ATSS) for short-range contacts. **D.** Distance distribution of ATSS-specific and shared PIRs at 5kb binned (baits unbinned) resolution. Top panel - interactions up to 1Mb (Kruskal-Wallis test p < 2.22e-16; pairwise Wilcoxon test p = 8.68e-6 [partially shared vs fully shared], p = 4.46e-8 [partially shared vs distinct] and p < 2.22e-16 [fully shared vs distinct]; bottom panel - interactions up to 50kb (Kruskal-Wallis test p = 7.65e-5; pairwise Wilcoxon test p = 9.8e-5 [partially shared vs fully shared], p = 6e-4 [partially shared vs distinct] and p = 1 [fully shared vs distinct]). **E.** Correlation between gene expression and number of regulatory elements identified in CHiCAGO PIRs at fragment and 5kb (solitary baits) resolution.

**Figure S2.**
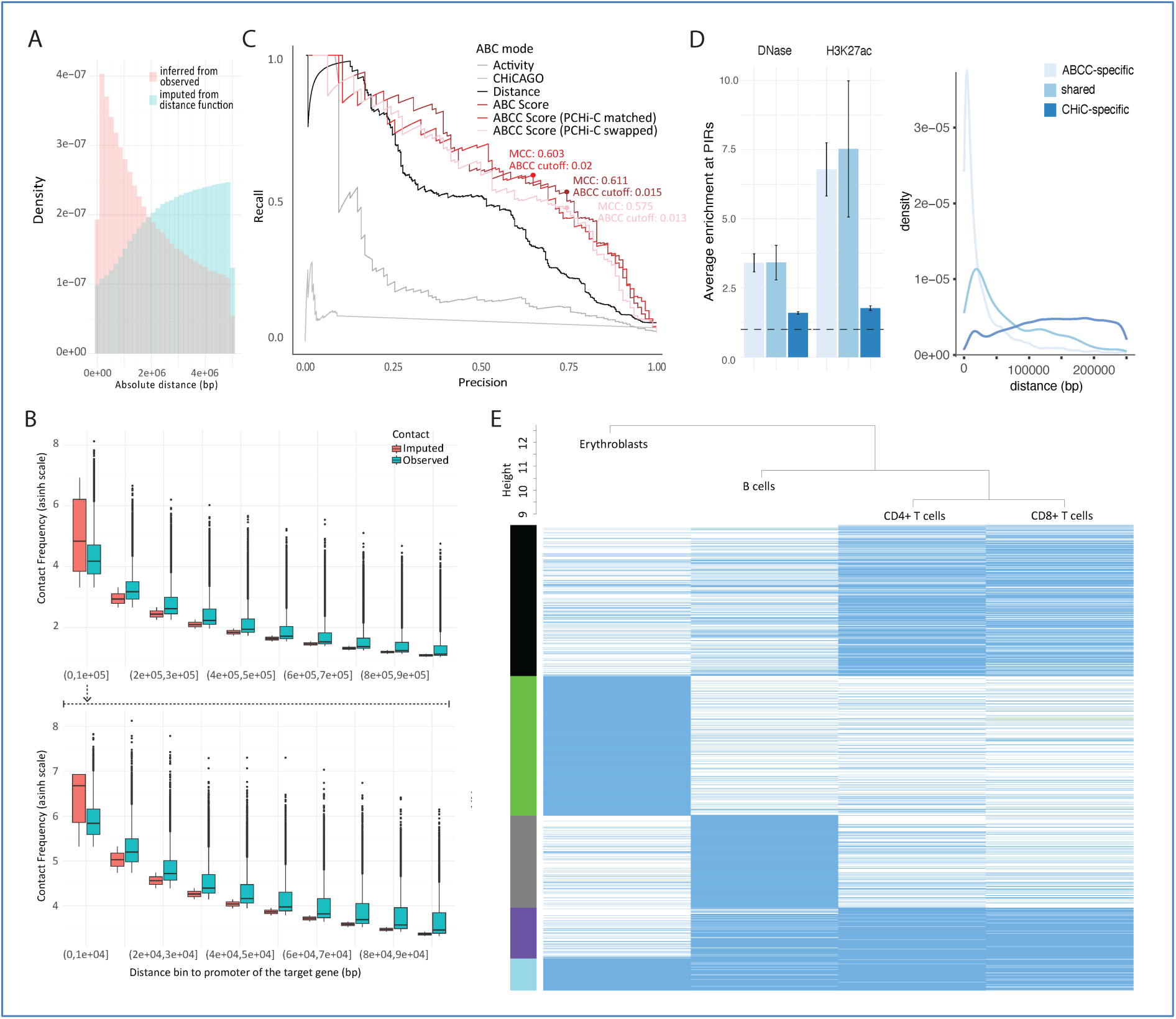
Benchmarking the ABCC approach with public data. **A.** Density distribution of promoter interactions inferred from observed PCHi-C contact frequencies (pink) and those imputed using the CHiCAGO distance function (cyan) across genomic interaction distances. **B.** Contact frequency distributions stratified by distance. Observed PCHi-C contacts are shown in green, imputed contacts (using expected frequencies estimated using the CHiCAGO distance function) are shown in blue. Similarly to standard ABC, frequency capping is introduced for short-range imputed contacts (<5kb). **C.** Precision–recall curves benchmarking the predictive performance of different scoring approaches for enhancer–promoter interactions in erythroblasts. Curves compare the scoring across: CHiCAGO-detected contacts, Activity alone, Distance alone, the conventional ABC score, and PCHi-C-based ABCC score in two modes: “matched” - using PCHi-C cell-type specific profile for erythroid cells and “swapped”, in which a PCHi-C dataset with a similar read coverage from a different cell type, CD4+ T cells, is used instead. MCC: Matthews correlation coefficient, an alternative to the AUC metric that is more informative under class imbalance and more sensitive to performance at a fixed decision threshold^175^. **D.** Enrichment of epigenetic markers at PIRs: DNase - chromatin accessibility and H3K27ac - active enhancers (left panel) and distance distribution of ABCC-specific, PCHiC-specific and shared enhancer-promoter links (right panel) in K562 cells for 0.023 ABCC threshold. **E.** Hierarchical clustering heatmap of enhancer–promoter interactions predicted with ABCC across cell types (erythroblasts, B cells, CD4+ T helper cells, CD8+ T cells).

**Figure S3.**
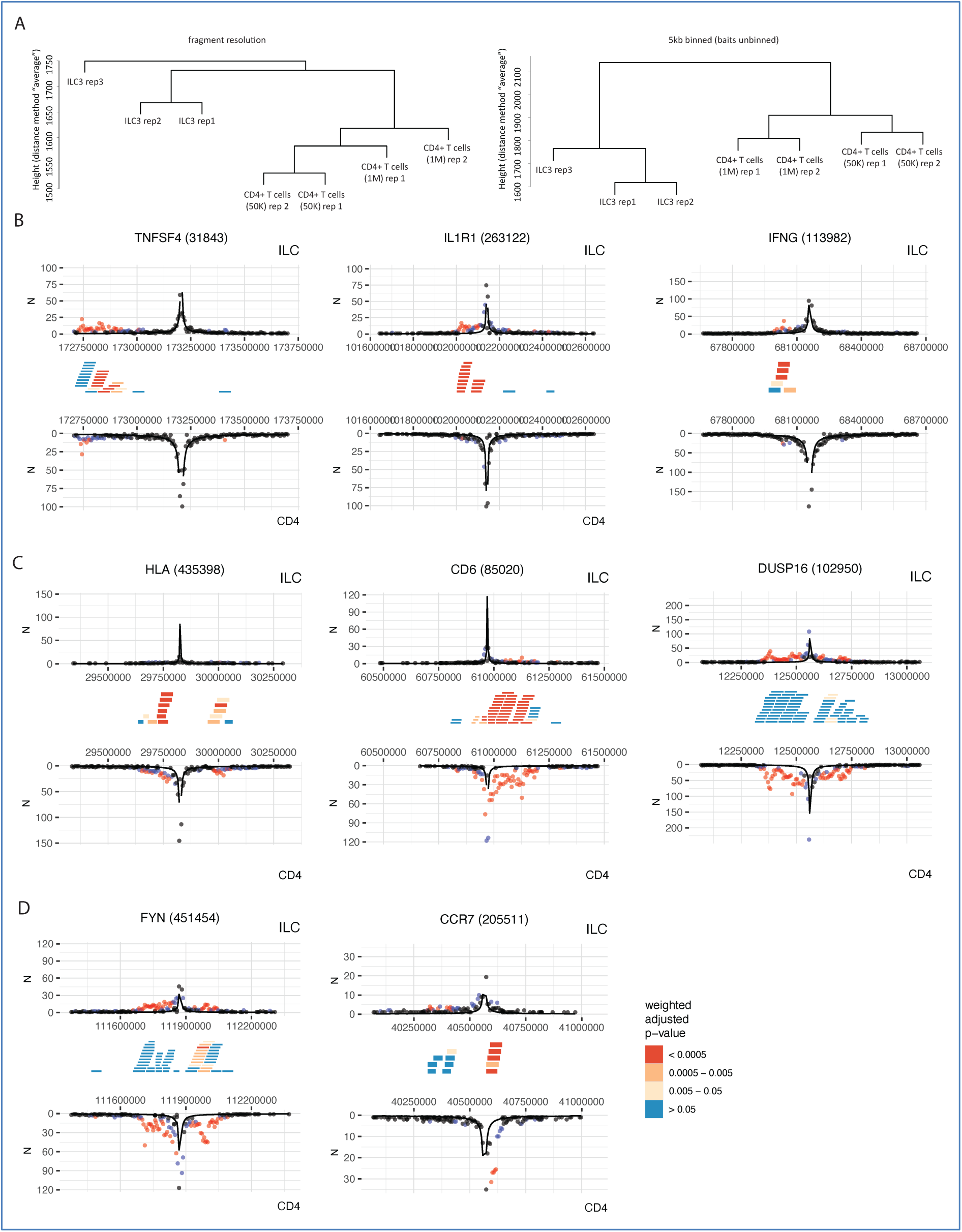
Genes with differential contacts in ILC3s and CD4+ T cells. **A.** Hierarchical clustering of ILC3s and CD4+ T cells PCHi-C datasets. **B-D.** Examples of captured promoters with differential wiring between ILC3s and CD4+ T cells: promoters with stronger **(B)** and weaker **(C)** contacts in ILC3s compared with CD4+ T cells, as well as with both types of contacts **(D).**

**Figure S4.**
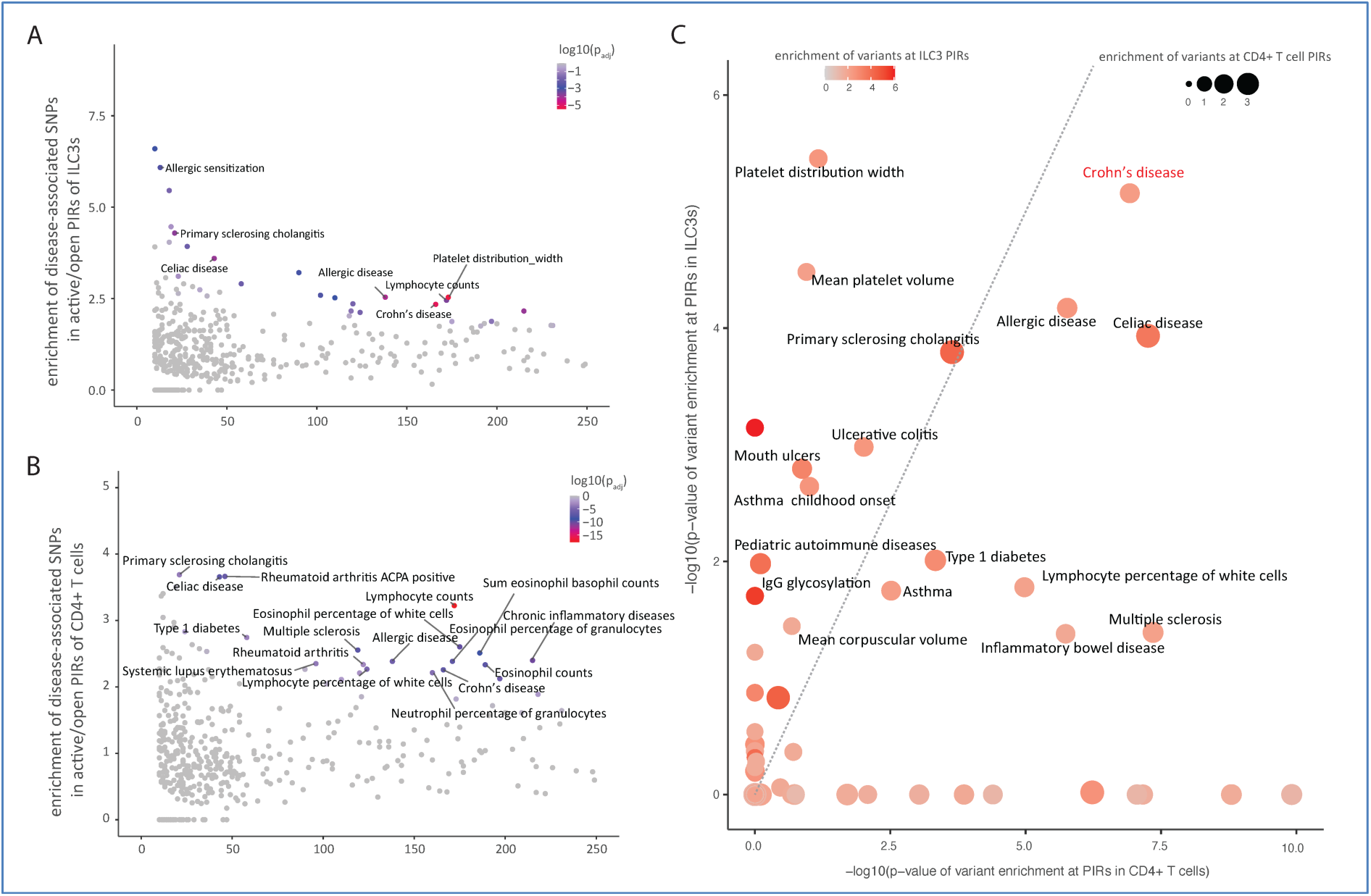
Supplementary information for the RELI analysis of risk loci enriched in ILC3 and CD4+ T-cell PIRs. **A-B** RELI enrichment of risk variants in ILC3s **(A)** and CD4s PIRs (**B**) across 495 diseases and traits. Traits with log_10_(BH corrected p-value in ILC3s) < 0.001, number of loci per trait > 10, and enrichment > 2.2 are labelled. **C.** Adjusted p-value of RELI enrichment of risk variants ILC3s vs CD4s PIRs across 495 diseases and traits. Traits with log_10_(BH corrected p-value in ILC3s) < 0.05 are labelled.

**Figure S5.**
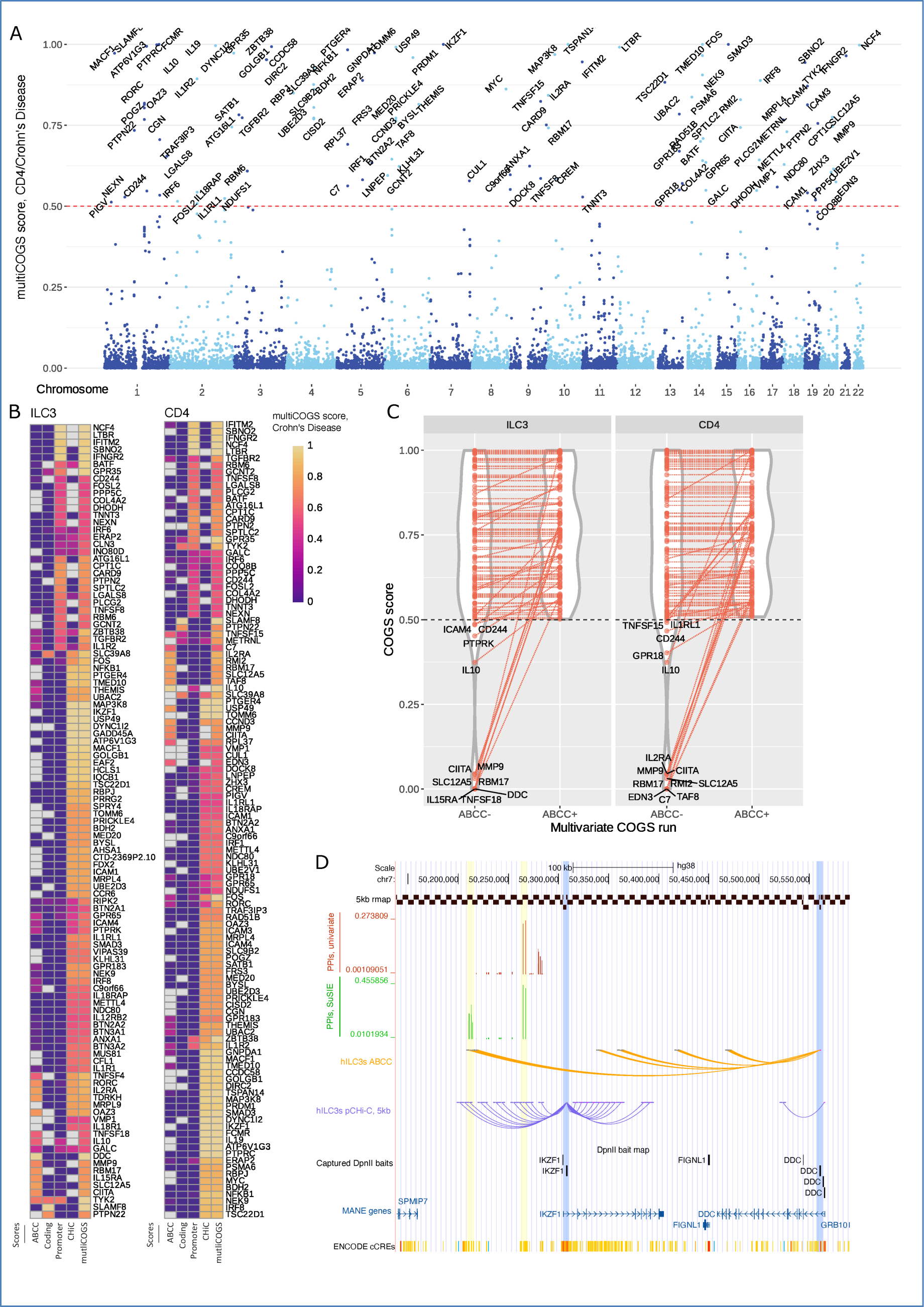
MultiCOGS prioritises gene sets in Crohn’s Disease. **A.** Manhattan plot showing multiCOGS for CD risk based on promoter contacts in CD4+ T cells. **B.** Heatmaps of region contributions to multiCOGS scores in ILC3s and CD4s in CD. **C.** Illustration of genes that were only prioritised for CD with the addition of ABCC, in ILC3s and CD4s. In each graph, the multiCOGS score with and without ABCC is plotted for all genes that were prioritised in the full multiCOGS run (score > 0.5 with ABCC). **D.** Illustration of multiCOGS prioritisation of *IKZF1* and *DDC* in ILC3s in the 7p locus. In this locus, multivariate fine mapping identifies two credible sets of variants (yellow bars), whereas univariate fine mapping only detects one. PCHi-C interactions connect these likely causal variants to the *IKZF1* promoter (first blue bar). However, ABCC interactions also connect one of the credible sets to the DDC promoter (second blue bar). Thus, multiCOGS prioritises both genes, whereas classic COGS prioritises only *IKZF1*.

**Figure S6.**
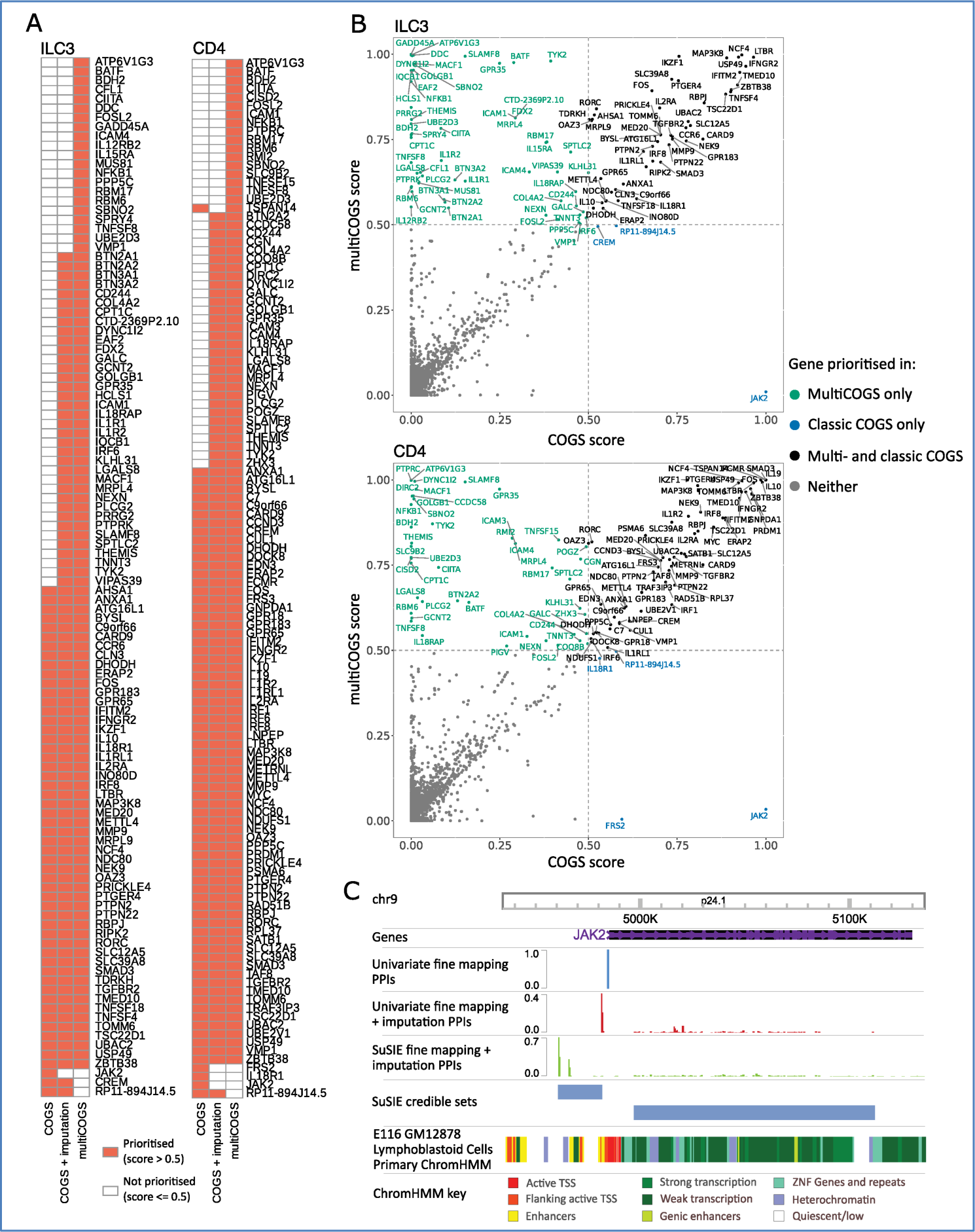
Comparison of gene prioritisation for Crohn’s Disease in classic COGS versus multiCOGS. **A.** Comparison of prioritised gene sets between classic COGS, classic COGS plus imputation, and multiCOGS (i.e. imputation plus multivariate fine mapping, processed via the multiCOGS algorithm) for CD. Shown for ILC3s and CD4+ T cells. **B.** Comparison of COGS scores and multiCOGS scores for genes in ILC3 cells (top) and CD4+ T cells (bottom) for CD. Green labels indicate genes prioritised in multiCOGS only, blue in classic COGS only, and black in both. **C.** Plot of the *JAK2* locus, showing the shift of the most likely causal variant from the promoter of *JAK2* to a region around 20kb upstream of the promoter upon multivariate fine mapping, leading to a lower multiCOGS vs classic COGS score. No chromosomal interactions were observed between this region and the *JAK2* promoter in ILC3 or CD4+ T cells.

**Figure S7.**
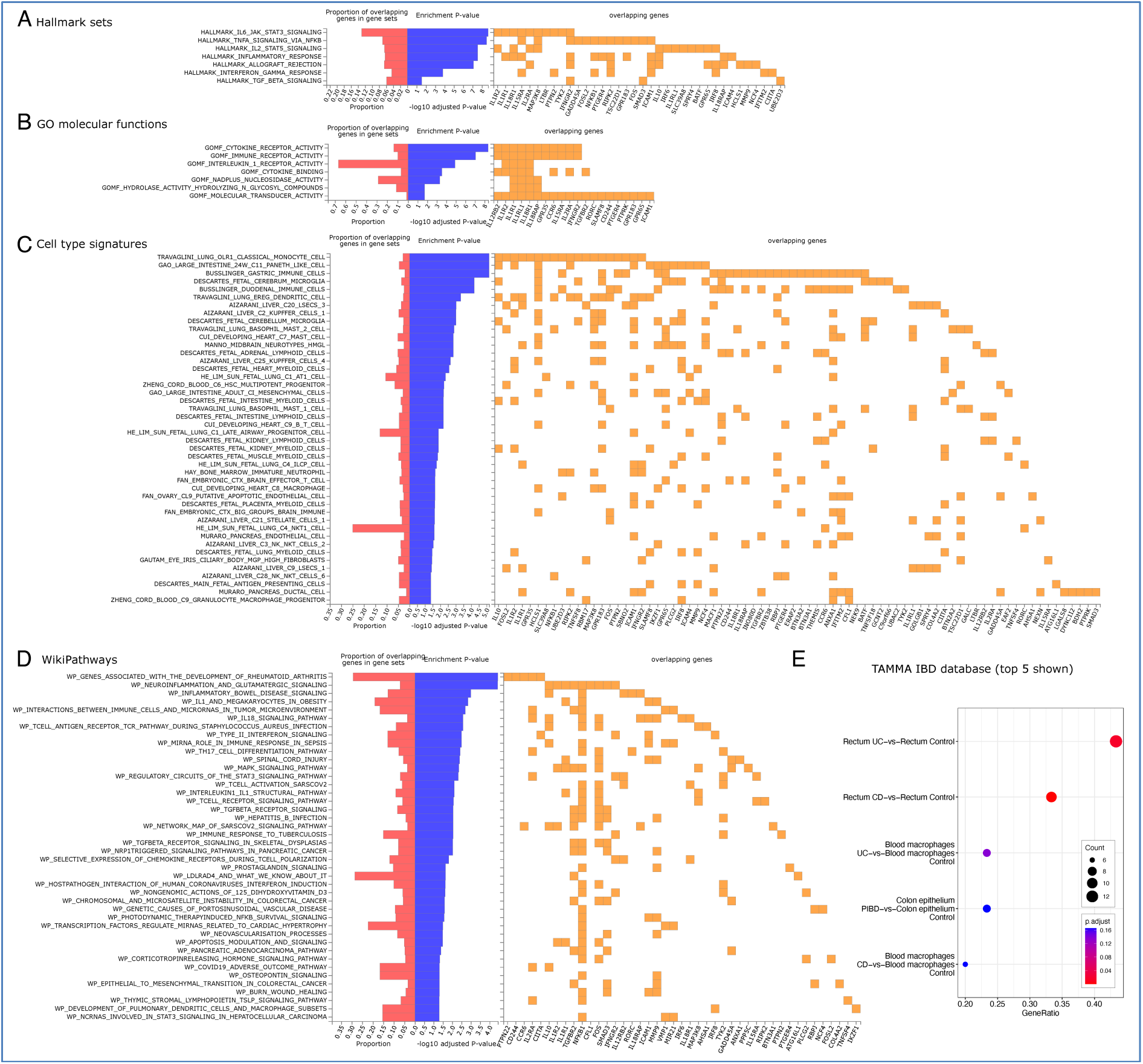
Biological annotation of multiCOGS CD genes in ILC3s. **A-D**. Enriched gene sets among multiCOGS genes detected using the GENE2FUNC pipeline in FUMA^121^, for the following databases: (**A**) MSigDB hallmark sets, (**B**) GO molecular functions, (**C**) MSigDB cell type signatures, (**D**) MSigDB WikiPathways. **E**. Enrichment analysis for differentially expressed gene sets among multiCOGS genes in the TAMMA IBD database.

**Figure S8.**
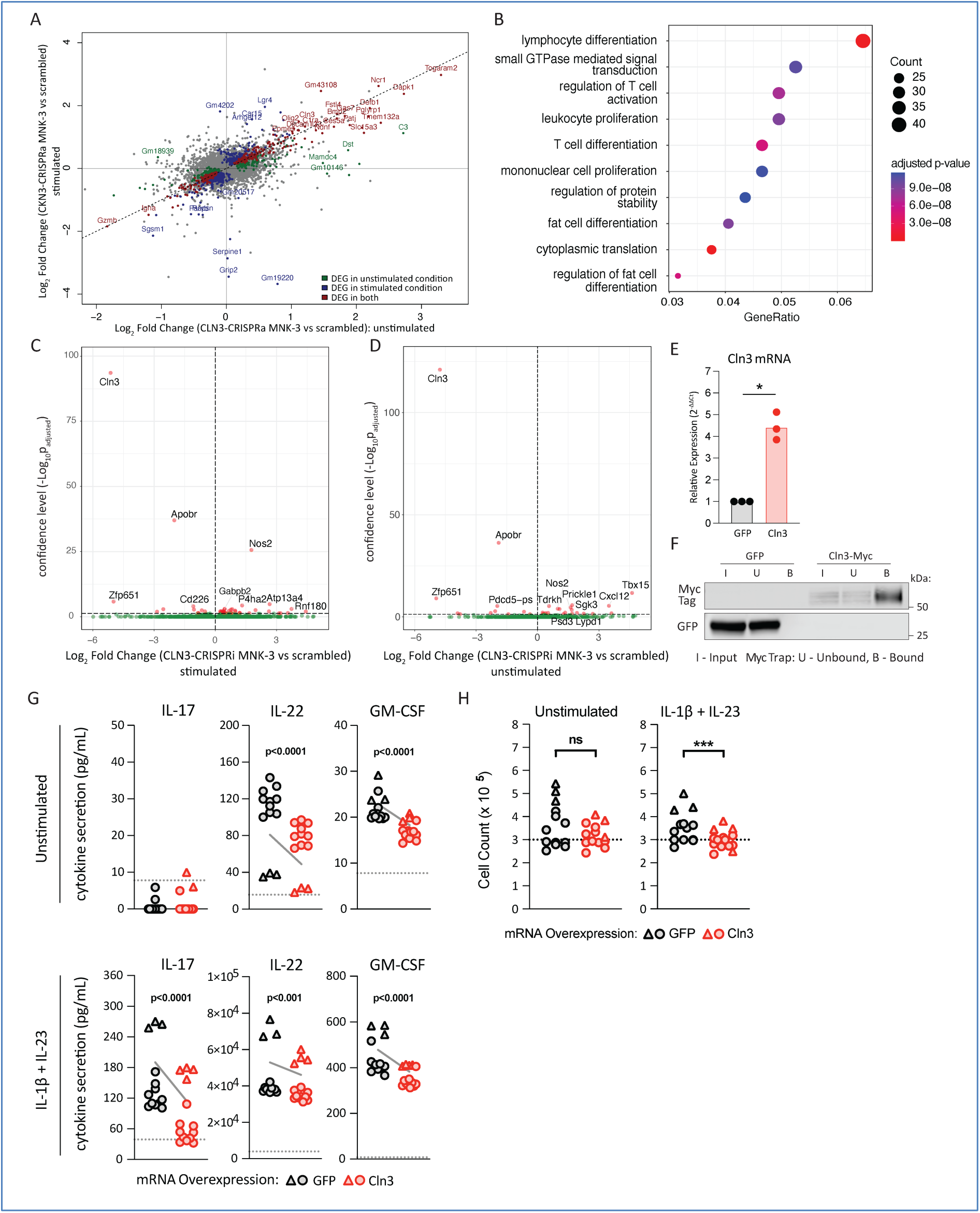
Additional information on the role of *Cln3* in ILC3 inflammatory function. **A.** Comparison of differentially expressed genes in IL23/IL-1βl-stimulated vs unstimulated *Cln3*-CRISPRa cells (relative to scrambled gRNA controls). **B.** GO term enrichment analysis for genes differentially expressed upon CLN3*-*CRISPRa stimulation. **C.** Differential expression of genes in IL23/IL-1βl-stimulated *Cln3*-CRISPRi MNK-3 cells vs scrambled gRNA controls. Red - differentially expressed genes (DESeq2 adjusted p-value < 0.05), green - all other genes. **D.** Differential expression of genes in unstimulated *Cln3*-CRISPRi MNK-3 cells vs scrambled gRNA controls. Red - differentially expressed genes (DESeq2 adjusted p-value < 0.05), green - all other genes. **E.** *Cln3* expression in MNK-3 cells electroporated with Cln3-myc mRNA or GFP mRNA. Transcript abundance was quantified by qPCR, normalised to *Hprt*, and expressed relative to the GFP mRNA control. Each point represents an independent experiment. Statistical significance was assessed using a paired Welch’s t-test, p<0.05 (*). **F.** Verification of CLN3-myc protein expression and Myc tag-dependent pulldown. MNK-3 cells were lysed, subjected to immunoprecipitation using Myc-Trap agarose, and resolved by reducing SDS-PAGE. “I” = input lysate; “U” = unbound fraction; “B” = bead-bound fraction. Immunoblotting with anti-myc tag antibody detected a ∼65–80 kDa Cln3-myc species selectively enriched in the bound fraction. **G.** Cytokine secretion upon *Cln3* overexpression across independent experiments. MNK-3 cells were electroporated with GFP mRNA (black) or Cln3-myc mRNA (red) and cultured for 24 hr under unstimulated (top row) or IL-1β + IL-23–stimulated (bottom row) conditions. Cytokine concentrations (IL-17, IL-22, GM-CSF) in culture supernatants were quantified by ELISA. Each symbol represents a biological replicate from two independent experiments (triangles vs circles). The dotted horizontal line indicates the lower limit of quantification for each assay. Statistical significance was assessed using a linear mixed-effects model with experiment as a random effect and transfection as a fixed effect (n=13–14 per condition). Solid grey lines indicate group means, with dotted grey bands indicating 95% confidence intervals of the fixed-effect. **H.** Cell numbers upon *Cln3* overexpression with and without inflammatory stimulation. MNK-3 cells were electroporated with GFP mRNA (black) or Cln3-myc mRNA (red) and cultured for 24 hr in unstimulated (left) or IL-1β + IL-23–stimulated (right) media. Each symbol represents a biological replicate from two independent experiments (triangles vs circles). Lines connect the experiment-specific means. Viable cell numbers were quantified by trypan blue exclusion. The dotted line indicates the number of cells seeded at 0 hr. Statistical significance was assessed using a linear mixed-effects model with experiment as a random effect and transfection as a fixed effect (n=13–14 per condition). Not significant (ns), p<0.001 (***).

**Figure S9.**
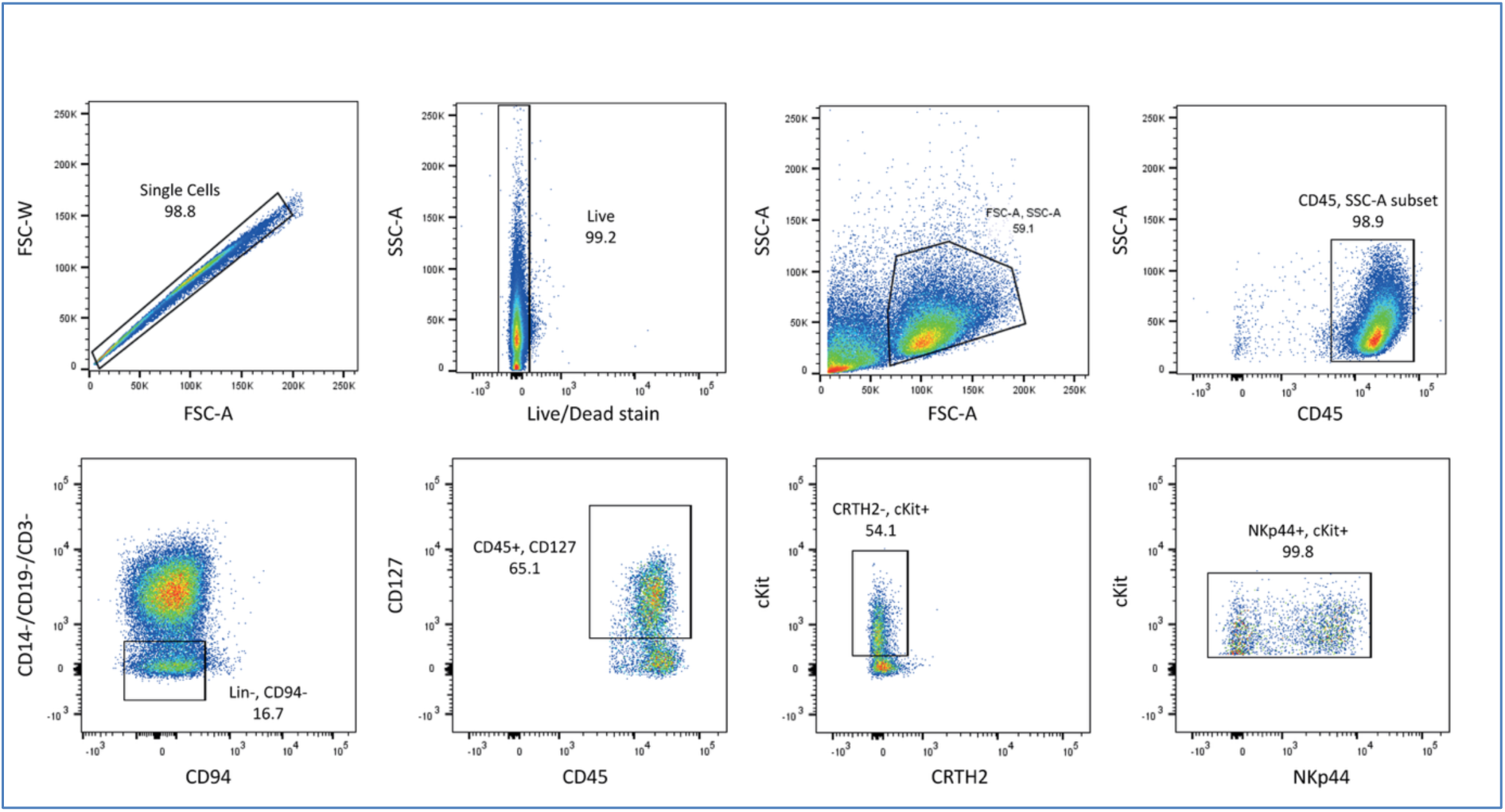
Flow cytometry gating strategy for isolation of human ILC3s from tonsils.

## List of supplementary tables

**Table S1.** PCHi-C quality metrics.

**Table S2.** Pathway enrichment for genes with ILC3-specific PIRs

**Table S3.** Pathway enrichment for genes with CD4-specific PIRs

**Table S4.** Pathway enrichment for genes with differential PIRs between ILC3 and CD4+ T cells

**Table S5.** Pathway enrichment for genes with non-differential PIRs between ILC3 and CD4+ T cells

**Table S6.** RELI results

**Table S7.** Candidate genes prioritised by multiCOGS in ILC3 cells and CD4+ T cells for Crohn’s Disease.

**Table S8.** Prior evidence for candidate genes prioritised by multiCOGS in ILC3 cells and CD4+ T cells for Crohn’s Disease.

**Table S9.** Pathway enrichment for multiCOGS-prioritised CD candidate genes in ILC3s

**Table S10.** LOLA results for TF enrichment within the PIRs of CD genes in ILC3s

**Table S11.** Differentially expressed genes upon CRISPR perturbations targeted to the *Cln3* promoter in MNK-3 cells

**Table S12.** MultiCOGS results in ILC3 and CD4+ T cells across 6 autoimmune traits

**Table S13.** The biological functions of multiCOGS-prioritised genes across 6 autoimmune traits in ILC3s

**Table S14.** sgRNA and primer sequences for Cln3 CRISPR targeting

## Supplementary Note 1

MultiCOGS resulted in loss of five candidate genes in one or both cell types, compared with classic COGS (*JAK2*, *CREM*, *FRS2*, *IL18R1* and *RP11-894J14.5*; see **Fig. S6B**). Of these, we were intrigued by the loss of *JAK2* in both cell types, because it is a well-noted candidate gene in IBD, with JAK inhibitors already used to treat ulcerative colitis and CD^172^. The COGS score for *JAK2* was substantially lower across both cell types when genetic imputation and multivariate fine mapping were employed (classic COGS score ∼1 in both cell types, multiCOGS score ∼0.01 in ILC3s and ∼0.03 in CD4s). Upon examining the locus, we discovered that fine mapping with the univariate methodology (Wakefield synthesis^173^) identified the most likely causal variant as rs1887428 (PPI = 0.999) at the *JAK2* promoter, but summary statistic imputation combined with multivariate fine mapping (SuSIE^69^) prioritised the variant rs1327500 (PPI = 0.663), in a region ∼20 kb upstream of *JAK2*, without detectable promoter contacts in ILC3 cells or CD4+ T cells (**Fig. S6C**). However, considering that both rs1887428 and rs1327500 are eQTLs for *JAK2* in blood cells, according to eQTLGen^174^, *JAK2* remains a strong candidate in this locus by genetic association.

